# Is amphistomy an adaptation to high light? Optimality models of stomatal traits along light gradients

**DOI:** 10.1101/601377

**Authors:** Christopher D. Muir

## Abstract

Stomata regulate the supply of CO_2_ for photosynthesis and the rate of water loss out of the leaf. The presence of stomata on both leaf surfaces, termed amphistomy, increases photosynthetic rate, is common in plants from high light habitats, and rare otherwise. In this study I use optimality models based on leaf energy budget and photosynthetic models to ask why amphistomy is common in high light habitats. I developed an R package **leafoptimizer** to solve for stomatal traits that optimally balance carbon gain with water loss in a given environment. The model predicts that amphistomy is common in high light because its marginal effect on carbon gain is greater than in the shade, but only if the costs of amphistomy are also lower under high light than in the shade. More generally, covariation between costs and benefits may explain why stomatal and other traits form discrete phenotypic clusters.

## Introduction

Stomata are microscopic pores formed by a pair of guard cells primarily located on the leaf surface of land plants. Their density and aperture on a leaf control the CO_2_ supply to leaf interiors and the rate of water lost through transpiration (recently reviewed in Sack & Buckley, 2016). Higher densities and/or larger pores allow more CO_2_ into the leaf, increasing photosynthetic rate, but also increasing transpiration (Farquhar & Sharkey, 1982). As the balance of CO_2_ and water demand and supply shifts through time and space, stomata respond over minutes to daily environmental variation, throughout the life of a single plant, and over long periods of evolutionary time (Wolfe, 1971; Woodward, 1987; Royer, 2001; Beerling & Royer, 2011; Milla *et al.*, 2013; McElwain & Steinthorsdottir, 2017).

A less appreciated aspect of stomata is that most leaves have all their stomata on the lower (usually abaxial) surface of the leaf, termed hypostomy, while some have them on both surfaces, termed amphistomy (Metcalfe & Chalk, 1950; Peat & Fitter, 1994; Muir, 2015; Drake *et al.*, 2019). Although amphistomy is rare in general, it is common among high light plants (Salisbury, 1927; Mott *et al.*, 1984; Peat & Fitter, 1994; Bucher *et al.*, 2017; Jordan *et al.*, 2014; Muir, 2018; Drake *et al.*, 2019). Why is amphistomy common in high light habitats but rare elsewhere? Amphistomy creates a second parallel pathway for CO_2_ diffusion into the leaf, which should increase photosynthesis especially when there is a lot of resistance to diffusion in the mesophyll (Parkhurst, 1978; Gutschick, 1984; Jones, 1985; Parkhurst & Mott, 1990). We might then expect amphistomy to be common, but it is not, implying some cost of amphistomy. Amphistomy also increases transpiration by forming a second boundary layer conductance for water transport (Foster & Smith, 1986, this study), but it is not clear if this tradeoff, or some other, explains variation in stomatal ratio. To evaluate these hypotheses and generate testable predictions, we need theory to predict how trait optima change across environments, both plastically and adaptively. These are classic evolutionary questions.

Stomata are also a fascinating and useful system for understanding phenotypic evolution. Land plants, like all major groups, can thrive in vastly different niches because of their diverse forms and functions, adaptations that evolved over millions of years. Less appreciated, but equally important in the study of phenotypic evolution, is that organisms occupy a small fraction of the feasible phenotypic space that could evolve in principle. This is true of stomata, as I will explain below. Why do some trait values rarely or never evolve? Three broad hypotheses explain why certain phenotypes can be rare or even absent from nature: 1) **Developmental inaccessibility** - a trait value is physically possible and would be favored by selection, but cannot evolve because the developmental system prevents the right genetic variation from arising; 2) **Rare environments** - a trait value is physically possible and would be favored by selection, but is rare because the environment that favors it is itself rare; and 3) **Selection** - a trait value is physically possible but is universally less fit than other trait values. Often, these hypotheses might be referred to as different phenotypic constraints (Arnold, 1992), but this terminology can be fraught with confusion and competing interpretations. In this paper, I focus on evaluating hypothesis 3, but address others throughout. Developmental inaccessibility could be important if mutations that initiate stomatal development on the upper leaf surface cause it to have the same stomatal density and size as the lower surface. This would make it easy to evolve amphistomy from hypostomy, but difficult to evolve different stomatal densities on each surface. It is also not hard to imagine that if there are discrete niches in the environment, then trait values should cluster around values best suited to those niches (Fig. 1D-F). It is more difficult to explain why trait values would cluster when the underlying environment is continuous because this implies that intermediate phenotypes are not favored in intermediate environments (Fig. 1G-I). This pattern would imply a nonlinear relationship between trait optima and environmental gradients.

**Figure 1:**
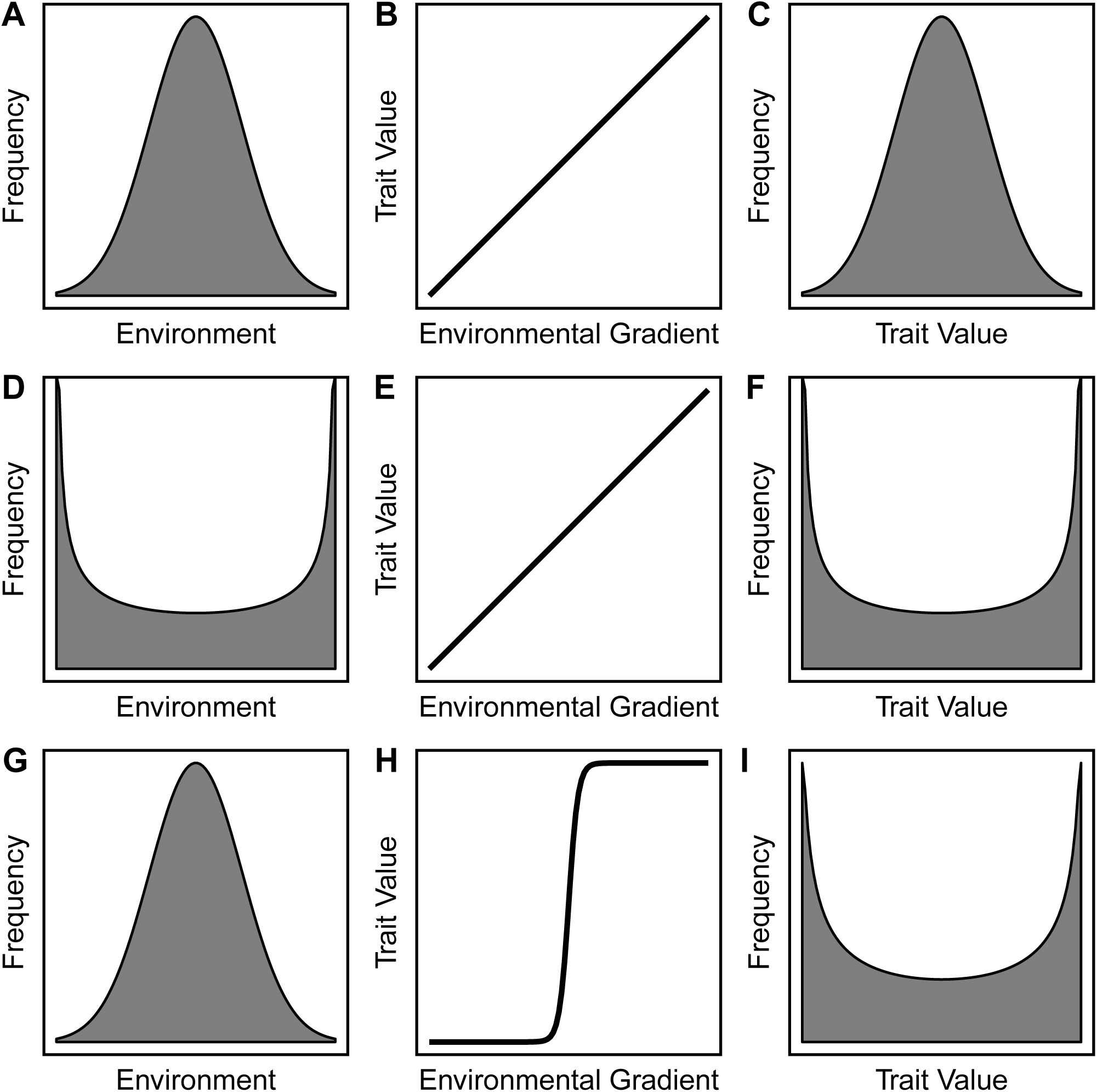
General hypotheses for clustering: No clustering occurs when the environment is unimodal and there is one-to-one matching between the environment and trait optima, leading to a unimodal trait distribution (top row, **A-C**). Clustering can occur if the environment is bimodal even with one-to-one matching between the environment and trait optima (middle row, **D-F**). Clustering can also occur if the environment is unimodal, but there is a nonlinear relationship between the environment and the trait optimum (bottom row, **G-I**). These latter two hypotheses are not mutually exclusive and may reinforce or counteract one another.

The ratio of stomatal densities on the upper surface to the sum of both surfaces (hereafter termed “stomatal ratio”), is a great system for studying why traits cluster because the distribution of this trait is highly clustered and we have mathematical tools to predict the optimal trait value in different environments. Stomatal ratio forms three main trait clusters in angiosperms (Muir, 2015): hypostomy (stomatal ratio = 0); complete amphistomy (stomatal ratio = 0.5); and hyperstomy (aka epistomy, stomatal ratio = 1). There are relatively few species with intermediate values, though they do exist and there is genetic variation, suggesting that development does not preclude the evolution of intermediate trait value (Muir *et al.*, 2014a,b, 2015). Few plants (mostly aquatic) are hyperstomatous (epistomatous), so I focus on the “bimodal” pattern describing two clusters, hypo- and amphistomy. Intermediate environments that favor intermediate stomatal ratios might be rare (Fig. 1D-F) or there may be a threshold-like relationship between the environment the trait optimum (Fig. 1H). To evaluate these hypotheses requires predictions about the relationship between the environment and trait optima.

Optimality models provide an independent way to predict the relationship between environments and trait optima against which we can compare observations of the natural world. They are an important part of identifying adaptive variation because”concordance between [optimality] model[s] and nature suggests adaptation” (Olson & Arroyo-Santos, 2015). Optimality models have a long history in successfully explaining plant form and function (Givnish, 1986, 1987), especially with stomata (Cowan & Farquhar, 1977; Buckley *et al.*, 2017b). Optimality models based on physics and chemistry are combined with a “goal” function to generate testable predictions about how traits *should* vary if organisms are adapted to their environment. If optimality models predict phenotypes that do not exist in nature, this might suggest developmental inaccessibility or rare environments prevent the phenotype from evolving. Optimality models may also fail if the “goal” function, assuming it is anything other than fitness, is misspecified.

In this study, I use optimality models to predict stomatal conductance and stomatal ratio across light gradients to evaluate under what conditions, if any, we would expect phenotypic clustering to evolve along a continuous environmental gradient (hypothesis 3). I also evaluate models on their ability to predict other, independent empirical observations. Ideally, a single model should account for all of the following observations: 1) amphistomy is rare (Metcalfe & Chalk, 1950; Peat & Fitter, 1994; Muir, 2015; Drake *et al.*, 2019); 2) amphistomy is more common in high light environments (Salisbury, 1927; Mott *et al.*, 1984; Mott & Michaelson, 1991; Peat & Fitter, 1994; Jordan *et al.*, 2014; Bucher *et al.*, 2017; Muir, 2018; Drake *et al.*, 2019); 3) amphistomy is associated with higher stomatal density (Beerling & Kelly, 1996; Muir, 2018), which is often a proxy for operational stomatal conductance (Franks & Beerling, 2009); and 4) stomatal ratio is bimodal (see above). Amphistomy is also more common in herbs than woody plants (Muir (2015, 2018), but see Drake *et al.* (2019)), but I do not address this observation here.

I examine three models with increasing complexity (Models 1–3). Model 1 assumes no extrinsic “cost” of amphistomy. It asks simply whether a tradeoff between carbon gain and water loss can explain the aforementioned empirical observations. Model 2 adds an extrinsic, *ad hoc* cost of amphistomy but is agnostic about the mechanism underlying this cost (see Discussion). Finally, Model 3 assumes that the extrinsic cost of amphistomy is not constant, but covaries with light gradients.

## Materials and Methods

I used biophysical and biochemical models of leaf temperature and photosynthesis to solve for the optimal stomatal conductance and stomatal ratio across different environments. The details of the leaf temperature and photosynthetic models are described elsewhere (Muir, 2019a,b), so I briefly summarize their structure here. A glossary of model inputs and outputs can be found in Tables 1 and 2, respectively. Values of photosynthetic temperature response functions and fixed parameters are described in Tables S1 and S2 – S4, respectively.

**Table 1:**
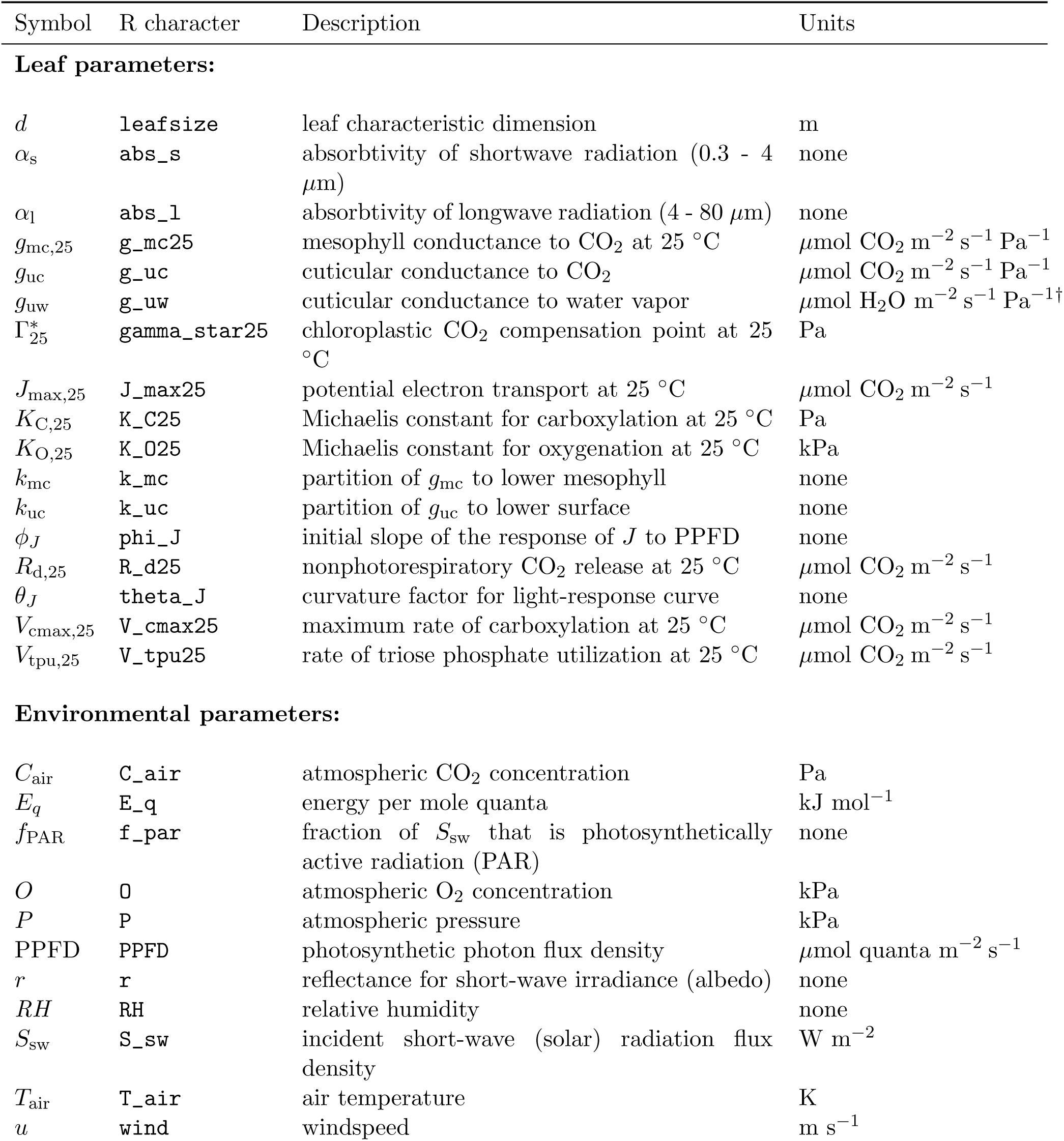
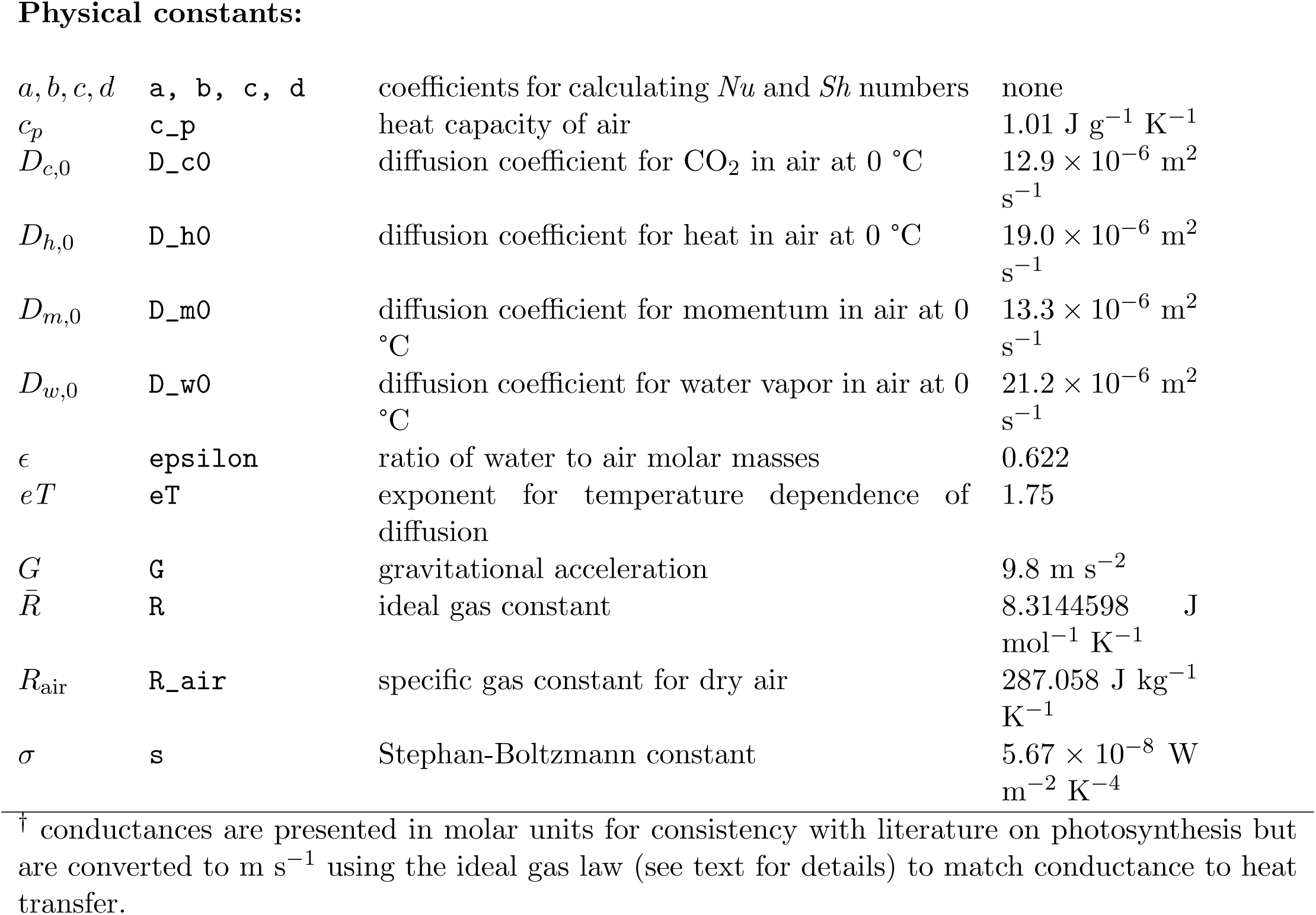
Parameter inputs for **leafoptimizer**. Each parameter has a mathematical symbol used in the text, the R character string used in the **leafoptimizer** package, a brief description, and the units. For physical constants, a value is provided where applicable, though users can modify these if desired. Conductances to CO_2_ (*g*_c_) are interconvertible with those for water vapour *g*_w_ and PPFD is interconvertible with *S*_sw_ (see Supporting Information).

| Symbol | R character | Description | Units |
| --- | --- | --- | --- |
| **Leaf parameters:** |  |  |  |
| $d$ | leafsize | leaf characteristic dimension | m |
| $\alpha_s$ | abs_s | absorbtivity of shortwave radiation (0.3 - 4 $\mu\text{m}$) | none |
| $\alpha_l$ | abs_l | absorbtivity of longwave radiation (4 - 80 $\mu\text{m}$) | none |
| $g_{mc,25}$ | g_mc25 | mesophyll conductance to CO<sub>2</sub> at 25 °C | $\mu\text{mol CO}_2 \text{ m}^{-2} \text{ s}^{-1} \text{ Pa}^{-1}$ |
| $g_{uc}$ | g_uc | cuticular conductance to CO<sub>2</sub> | $\mu\text{mol CO}_2 \text{ m}^{-2} \text{ s}^{-1} \text{ Pa}^{-1}$ |
| $g_{uw}$ | g_uw | cuticular conductance to water vapor | $\mu\text{mol H}_2\text{O m}^{-2} \text{ s}^{-1} \text{ Pa}^{-1}\dagger$ |
| $\Gamma_{25}^*$ | gamma_star25 | chloroplastic CO<sub>2</sub> compensation point at 25 °C | Pa |
| $J_{\text{max},25}$ | J_max25 | potential electron transport at 25 °C | $\mu\text{mol CO}_2 \text{ m}^{-2} \text{ s}^{-1}$ |
| $K_{C,25}$ | K_C25 | Michaelis constant for carboxylation at 25 °C | Pa |
| $K_{O,25}$ | K_O25 | Michaelis constant for oxygenation at 25 °C | kPa |
| $k_{mc}$ | k_mc | partition of $g_{mc}$ to lower mesophyll | none |
| $k_{uc}$ | k_uc | partition of $g_{uc}$ to lower surface | none |
| $\phi_J$ | phi_J | initial slope of the response of $J$ to PPFD | none |
| $R_{d,25}$ | R_d25 | nonphotorespiratory CO<sub>2</sub> release at 25 °C | $\mu\text{mol CO}_2 \text{ m}^{-2} \text{ s}^{-1}$ |
| $\theta_J$ | theta_J | curvature factor for light-response curve | none |
| $V_{\text{cmax},25}$ | V_cmax25 | maximum rate of carboxylation at 25 °C | $\mu\text{mol CO}_2 \text{ m}^{-2} \text{ s}^{-1}$ |
| $V_{\text{tpu},25}$ | V_tpu25 | rate of triose phosphate utilization at 25 °C | $\mu\text{mol CO}_2 \text{ m}^{-2} \text{ s}^{-1}$ |
| **Environmental parameters:** |  |  |  |
| $C_{\text{air}}$ | C_air | atmospheric CO<sub>2</sub> concentration | Pa |
| $E_q$ | E_q | energy per mole quanta | $\text{kJ mol}^{-1}$ |
| $f_{\text{PAR}}$ | f_par | fraction of $S_{sw}$ that is photosynthetically active radiation (PAR) | none |
| $O$ | O | atmospheric O<sub>2</sub> concentration | kPa |
| $P$ | P | atmospheric pressure | kPa |
| PPFD | PPFD | photosynthetic photon flux density | $\mu\text{mol quanta m}^{-2} \text{ s}^{-1}$ |
| $r$ | r | reflectance for short-wave irradiance (albedo) | none |
| $RH$ | RH | relative humidity | none |
| $S_{sw}$ | S_sw | incident short-wave (solar) radiation flux density | $\text{W m}^{-2}$ |
| $T_{\text{air}}$ | T_air | air temperature | K |
| $u$ | wind | windspeed | $\text{m s}^{-1}$ |
| **Physical constants:** |  |  |  |
| $a, b, c, d$ | **a, b, c, d** | coefficients for calculating $Nu$ and $Sh$ numbers | none |
| $c_p$ | **c_p** | heat capacity of air | $1.01 \text{ J g}^{-1} \text{ K}^{-1}$ |
| $D_{c,0}$ | **D_c0** | diffusion coefficient for $\text{CO}_2$ in air at $0^\circ\text{C}$ | $12.9 \times 10^{-6} \text{ m}^2 \text{ s}^{-1}$ |
| $D_{h,0}$ | **D_h0** | diffusion coefficient for heat in air at $0^\circ\text{C}$ | $19.0 \times 10^{-6} \text{ m}^2 \text{ s}^{-1}$ |
| $D_{m,0}$ | **D_m0** | diffusion coefficient for momentum in air at $0^\circ\text{C}$ | $13.3 \times 10^{-6} \text{ m}^2 \text{ s}^{-1}$ |
| $D_{w,0}$ | **D_w0** | diffusion coefficient for water vapor in air at $0^\circ\text{C}$ | $21.2 \times 10^{-6} \text{ m}^2 \text{ s}^{-1}$ |
| $\epsilon$ | **epsilon** | ratio of water to air molar masses | 0.622 |
| $eT$ | **eT** | exponent for temperature dependence of diffusion | 1.75 |
| $G$ | **G** | gravitational acceleration | $9.8 \text{ m s}^{-2}$ |
| $\bar{R}$ | **R** | ideal gas constant | $8.3144598 \text{ J mol}^{-1} \text{ K}^{-1}$ |
| $R_{\text{air}}$ | **R_air** | specific gas constant for dry air | $287.058 \text{ J kg}^{-1} \text{ K}^{-1}$ |
| $\sigma$ | **s** | Stephan-Boltzmann constant | $5.67 \times 10^{-8} \text{ W m}^{-2} \text{ K}^{-4}$ |

<sup>†</sup> conductances are presented in molar units for consistency with literature on photosynthesis but are converted to $\text{m s}^{-1}$ using the ideal gas law (see text for details) to match conductance to heat transfer.

**Table 2:** Calculated parameter outputs for **leafoptimizer**. Each parameter has a mathematical symbol used in the text, the R character string used in the **leafoptimizer** package, a brief description, and the units. Note that *g*_sc,opt_ is interconvertible with *g*_sw,opt_ and *k*_sc,opt_ is interconvertible with *SR*_opt_ (see Supporting Information).

| Symbol | R character | Description | Units |
| --- | --- | --- | --- |
| **Optimized leaf parameters:** |  |  |  |
| $g_{sc,opt}$ | `g_sc` | optimal stomatal conductance to CO<sub>2</sub> | $\mu\text{mol CO}_2 \text{ m}^{-2} \text{ s}^{-1} \text{ Pa}^{-1}$ |
| $g_{sw,opt}$ | `g_sw` | optimal stomatal conductance to water vapor | $\mu\text{mol H}_2\text{O m}^{-2} \text{ s}^{-1} \text{ Pa}^{-1}$ |
| $k_{sc,opt}$ | `k_sc` | optimal partition of $g_{sc,opt}$ to lower surface | none |
| $SR_{opt}$ | `sr` | optimal stomatal ratio | none |
| **Leaf parameters:** |  |  |  |
| $A$ | `A` | photosynthetic rate | $\mu\text{mol CO}_2 \text{ m}^{-2} \text{ s}^{-1}$ |
| $C_{chl}$ | `C_chl` | chloroplastic CO<sub>2</sub> concentration | Pa |
| $E$ | `E` | transpiration rate | $\text{mol H}_2\text{O m}^{-2} \text{ s}^{-1}$ |
| $g_h$ | `g_h` | boundary layer conductance to heat | $\text{m s}^{-1}$ |
| $g_{bc}$ | `g_bc` | boundary layer conductance to CO<sub>2</sub> | $\mu\text{mol CO}_2 \text{ m}^{-2} \text{ s}^{-1} \text{ Pa}^{-1}$ |
| $g_{bw}$ | `g_bw` | boundary layer conductance to water vapor | $\mu\text{mol H}_2\text{O m}^{-2} \text{ s}^{-1} \text{ Pa}^{-1}$ |
| $g_{mc}$ | `g_mc` | mesophyll conductance to CO<sub>2</sub> at $T_{leaf}$ | $\mu\text{mol CO}_2 \text{ m}^{-2} \text{ s}^{-1} \text{ Pa}^{-1}$ |
| $g_{tc}$ | `g_tc` | total conductance to CO<sub>2</sub> | $\mu\text{mol CO}_2 \text{ m}^{-2} \text{ s}^{-1} \text{ Pa}^{-1}$ |
| $g_{tw}$ | `g_tw` | total conductance to water vapor | $\mu\text{mol H}_2\text{O m}^{-2} \text{ s}^{-1} \text{ Pa}^{-1}$ |
| $\Gamma^*$ | `gamma_star` | chloroplastic CO<sub>2</sub> compensation point at $T_{leaf}$ | Pa |
| $Gr$ | `Gr` | Grashof number | none |
| $H$ | `H` | sensible heat flux density | $\text{W m}^{-2}$ |
| $J_{max}$ | `J_max` | potential electron transport at $T_{leaf}$ | $\mu\text{mol CO}_2 \text{ m}^{-2} \text{ s}^{-1}$ |
| $K_C$ | `K_C` | Michaelis constant for carboxylation at $T_{leaf}$ | Pa |
| $K_O$ | `K_O` | Michaelis constant for oxygenation at $T_{leaf}$ | kPa |
| $L$ | `L` | latent heat flux density | $\text{W m}^{-2}$ |
| $Nu$ | `Nu` | Nusselt number | none |
| $R_{abs}$ | `R_abs` | total absorbed radiation | $\text{W m}^{-2}$ |
| $R_d$ | `R_d` | nonphotorespiratory CO<sub>2</sub> release at $T_{leaf}$ | $\mu\text{mol CO}_2 \text{ m}^{-2} \text{ s}^{-1}$ |
| $Re$ | `Re` | Reynolds number | none |
| $Sh$ | `Sh` | Sherwood number | none |
| $S_r$ | `S_r` | longwave re-radiation | $\text{W m}^{-2}$ |
| $T_{leaf}$ | `T_leaf` | leaf temperature | K |
| $V_{cmax}$ | `V_cmax` | maximum rate of carboxylation at $T_{leaf}$ | $\mu\text{mol CO}_2 \text{ m}^{-2} \text{ s}^{-1}$ |
| $V_{tpu}$ | `V_tpu` | rate of triose phosphate utilization at $T_{leaf}$ | $\mu\text{mol CO}_2 \text{ m}^{-2} \text{ s}^{-1}$ |
| **Temperature-dependent physical parameters:** |  |  |  |
| $D_c$ | `D_c` | diffusion coefficient for CO<sub>2</sub> in air at $T_{leaf}$ | $\text{m}^2 \text{ s}^{-1}$ |
| $D_h$ | `D_h` | diffusion coefficient for heat in air at $T_{leaf}$ | $\text{m}^2 \text{ s}^{-1}$ |
| $D_m$ | `D_m` | diffusion coefficient for momentum in air at $T_{leaf}$ | $\text{m}^2 \text{ s}^{-1}$ |
| $D_w$ | `D_w` | diffusion coefficient for water vapor in air at $T_{leaf}$ | $\text{m}^2 \text{ s}^{-1}$ |

### Leaf temperature model

I modeled equilibrium leaf temperature using energy budget models (recently reviewed in Gutschick, 2016) implemented using the R package **tealeaves** version 1.0.1 (Muir, 2019b). Given a set of leaf parameters, environmental parameters, and physical constants, leaf energy budget models find the leaf temperature (*T*_leaf_) such that the net energy flux in W m^−2^ is balanced:

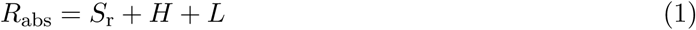

where *R*_abs_ is the absorbed radiation, *S*_r_ is infrared re-radiation, *H* is sensible heat loss, and *L* is latent heat loss. Absorbed radiation and infrared re-radiation are largely determined by the environment and not affected by stomatal traits. Leaf traits, especially leaf size, strongly impact sensible heat flux (*H*), but stomatal traits do not affect these properties directly. Stomatal traits strongly affect the total conductance to water vapor (*g*_tw_), which is proportional to the latent heat lost (*L*) as liquid water vaporizes and exits the leaf as a gas:

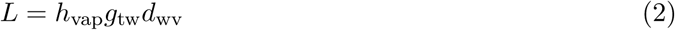

**tealeaves** models the latent heat of vaporization (*h*_vap_) as a linear function of temperature (Muir, 2019b; Nobel, 2009). *d*_wv_ is the water vapor pressure differential from the inside to the outside of leaf in units of mol m^−3^:

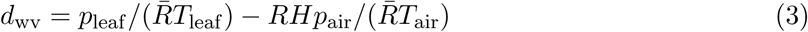

I assume the leaf interior is fully saturated (*p*_leaf_ is the saturation water vapor pressure at *T*_leaf_), where the saturation water vapor pressure is a function of temperature calculated using the Goff-Gratch equation (Vömdel, 2016). The vapor pressure of air is the product of the relative humidity (*RH*) and *p*_air_, the saturation water vapor pressure *T*_air_.

*g*_tw_ is the sum of the parallel lower (usually abaxial) and upper (usually adaxial) conductances in units of m s^−1^, which is the convention in leaf energy budget models:

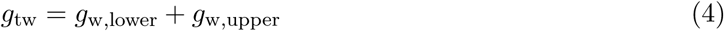

The conductance to water vapor on each surface (indexed as *j*) is a function of parallel stomatal (*g*_sw,*j*_) and cuticular (*g*_uw,*j*_) conductances in series with the boundary layer conductance (*g*_bw,*j*_). The stomatal and cuticular conductances on the lower surface are:

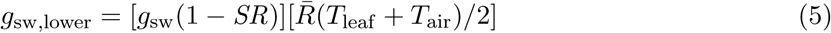

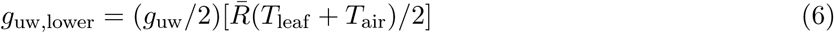

Note that the *total* leaf stomatal and cuticular conductances (*g*_sw_ and *g*_uw_ respectively) are in units of *µ*mol H_2_O m^−2^ s^−1^ Pa^−1^ in keeping with conventions of photosynthetic models (see below). In the above equations, these values are converted to units of m s^−1^ using the ideal gas law for the leaf energy budget model. Stomatal conductance is partitioned among leaf surfaces depending on stomatal ratio (*SR*). When *SR* = 0, all conductance is on the lower surface; when *SR* = 1, all conductance is on the upper surface; when *SR* = 0.5, conductance is evenly divided across surfaces. Cuticular conductance is assumed equal on each leaf surface, though this is probably not true in nature (Karbulková *et al.*, 2008). The corresponding expressions for the upper surface are:

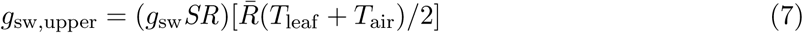

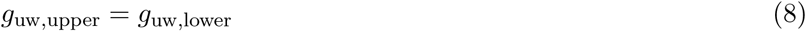

The boundary layer conductances for each surface differ because free convection differs on each surface (Foster & Smith, 1986):

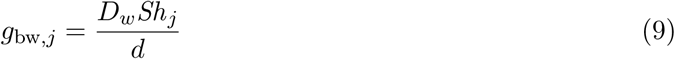

*d* is the leaf characteristic dimension in m, a physiologically relevant measure of leaf size because it determines heat and mass transfer (Taylor, 1975; Leigh *et al.*, 2017). *D*_*w*_ is the diffusion coefficient of water vapor in air as a function of temperature in units of m^2^ s^−1^:

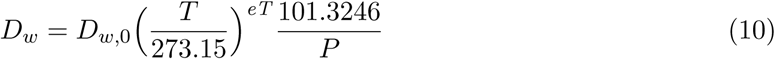

Each surface has its own unitless Sherwood number (*Sh*) that is a mix of free and forced convection:

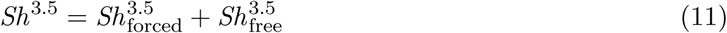

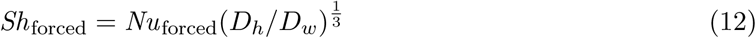

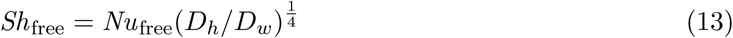

The Nusselt number (*Nu*) is a dimensionless number for heat transfer (Monteith & Unsworth, 2013). Free convection dominates when the Archimedes number (*Ar*) is greater than 10; forced convection dominates when *Ar* ≪ 0.1 (Nobel, 2009). Forced convection is probably most common in nature (Jones, 2014), but free convection can be important for large leaves at low wind speeds (see Muir, 2019b, for further detail). Because free convection depends on gravity, horizontally oriented leaves will exchange latent heat differently depending on how transpiration through stomata is distributed between surfaces. *D*_*h*_ is the diffusion coefficient of heat in air a function of temperature in units of m^2^ s^−1^, calculated following Eqn 10 with *D*_*h*,0_ substituted for *D*_*w*,0_ (Table 1).

Transpiration rate (mol H_2_O m^−2^ s^−1^) is the product of the total conductance to water vapour (Eqn 4) and the water vapor gradient (Eqn 3):

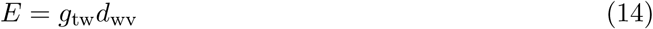

Foster & Smith (1986) previously demonstrated that amphistomatous leaves transpire more water than hypostomatous leaves at low wind speeds, holding total *g*_sw_ constant. To illustrate this result, I analyzed a similar model using **tealeaves** for hypostomatous (*SR* = 0), intermediate (*SR* = 0.25), and amphistomatous (*SR* = 0.5) leaves. I varied wind speed between 0 and 2 m s^−1^ at two light levels, photosynthetic photon flux density (PPFD) = 500 (shade) and 1500 (sun) *µ*mol quanta m^−2^ s^−1^. I fixed other leaf parameters as absortivity of shortwave radiation (*α*_s_) = 0.5, absorbtivity of longwave radiation (*α*_l_) = 0.97, *d* = 0.1 m, *g*_sw_ = 2 *µ*mol H_2_O m^−2^ s^−1^ Pa^−1^, *g*_uw_ = 0.1 *µ*mol H_2_O m^−2^ s^−1^ Pa^−1^. I fixed other environmental parameters where: atmospheric pressure (*P*) = 101.3246 kPa, relative humidity (*RH*) = 0.5, albedo (*r*) = 0.2, and air temperature (*T*_air_) = 25 °C. Physical constants are described in Table 1. I calculated the ratio of transpiration for an intermediate or amphistomatous leaf (*E*_*j*_) compared to that of hypotstomatous (*E*_hypo_) leaf in the same environment:

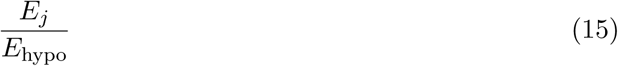

### Photosynthesis model

The **photosynthesis** package version 1.0.1 (Muir, 2019a) implements the Farquhar-von Caemmerer-Berry biochemical model of C_3_ photosynthesis (Farquhar *et al.*, 1980), which has been reviewed extensively elsewhere (e.g. Sharkey *et al.*, 2007). Following the treatment of Buckley & Diaz-Espejo (2015), the photosynthetic demand rate (*A*_*D*_) is the minimum of Rubisco-, RuBP regeneration-, and TPU-limited assimilation rates:

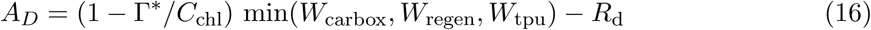

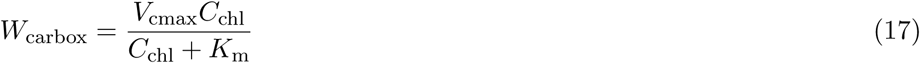

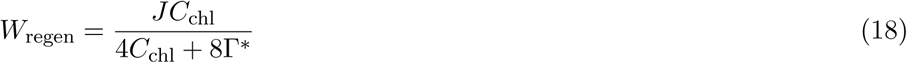

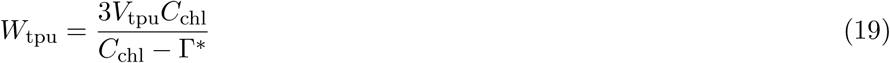

*K*_m_ is the Michaelis-Menten constant:

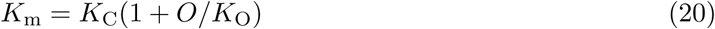

*J* is a function Photosynthetic photon flux density (PPFD), obtained by solving the equation:

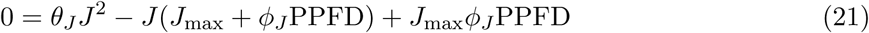

The photosynthetic supply rate (*A*_*S*_) is the product of the total conductance to CO_2_ (*g*_tc_ [*µ*mol CO_2_ m^−2^ s^−1^ Pa^−1^]) and CO_2_ drawdown (*C*_air_ − *C*_chl_):

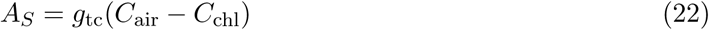

To facilitate modeling differentiated upper and lower leaf anatomies, **photosynthesis** allows users to partition boundary, cuticular, stomatal, and mesophyll conductances separately to each surface (similar to Jones, 1985). On surface *j*, there are two parallel conductances, the cuticular conductance (*g*_uc,*j*_) and the in-series conductances through mesophyll (*g*_mc,*j*_), stomata (*g*_sc,*j*_), and boundary layer (*g*_bc,*j*_). Following rules for circuits (Nobel, 2009), the total conductance for surface *j* is:

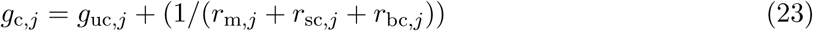

To simplify the formula, I substitute resistance for conductance 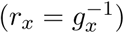 above. Boundary layer conductances to CO_2_ are calculated as described above for water vapor, but accounting for the different diffusivity of CO_2_ and water vapor in air (see Supporting Information for detail). The mesophyll (*g*_mc_) conductance is partitioned between layers using the following definitions:

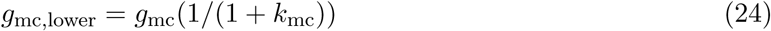

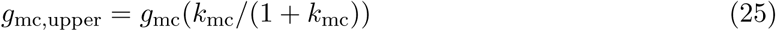

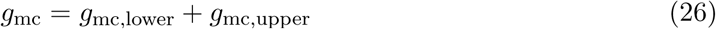

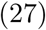

*g*_mc_ is the total leaf conductance through meosphyll, partitioned to lower or upper leaf portions based on the partitioning factor *k*_mc_. The cuticular conductance to CO_2_ (*g*_uc_) is converted from that for water vapor (*g*_uw_, see Eqns 6, 8) as described in the Supporting Information.

I modeled photosynthetic temperature responses following Bernacchi *et al.* (2002) and Buckley & Diaz-Espejo (2015). Values of temperature-dependent parameters are provided at 25 °C as input (Table 1) and computed at *T*_leaf_ (Table 2) to determine the photosynthetic rate. The photosynthetic rate *A* at a given *T*_leaf_ is determined by solving for the *C*_chl_ that balances photosynthetic supply and demand rates (*A*_*D*_ = *A*_*S*_).

Parkhurst (1978), Gutschick (1984), and Jones (1985) previously demonstrated that amphistomatous leaves should photosynthesize more than hypostomatous leaves holding other factors constant. To illustrate this result, I used the **photosynthesis** package to model photosynthetic rate for hypostomatous (*SR* = 0), intermediate (*SR* = 0.25), and amphistomatous (*SR* = 0.5) leaves. I varied *T*_leaf_ between 5 and 40 °C at two levels of *g*_sw_, 1 (low) and 4 (high) *µ*mol H_2_O m^−2^ s^−1^ Pa^−1^. I fixed other leaf parameters as *g*_mc,25_ = 3 *µ*mol CO_2_ m^−2^ s^−1^ Pa^−1^, *g*_uc_ = 0.1 *µ*mol CO_2_ m^−2^ s^−1^ Pa^−1^, *d* = 0.1 m, *J*_max,25_ = 150 *µ*mol CO_2_ m^−2^ s^−1^, *ϕ*_*J*_ = 0.331, *R*_d,25_ = 2 *µ*mol CO_2_ m^−2^ s^−1^, *θ*_*J*_ = 0.825, *V*_cmax,25_ = 100 *µ*mol CO_2_ m^−2^ s^−1^, *V*_tpu,25_ = 200 *µ*mol CO_2_ m^−2^ s^−1^. I fixed other environmental parameters where: *C*_air_ = 41 Pa, *O* = 21.27565 kPa, *P* = 101.3246 kPa, PPFD = 1500 *µ*mol quanta m^−2^ s^−1^, *RH* = 0.5, *T*_air_ = *T*_leaf_, and *u* = 2 m s^−1^. Physical constants are described in Table 1.

### Optimization of stomatal traits

Biophysical and biochemical models like those implemented in **tealeaves** and **photosynthesis** help understand structure-function relationships, but cannot by themselves predict ecological and evolutionary variation. Optimality models with a defined “goal” function make testable predictions about ecological and evolutionary responses to the environment (Givnish, 1986). In plant physiology, optimality models often assume that plants will modify stomatal traits through acclimation (within generations) or adaptation (between generations) to maximize carbon gain minus costs (usually water loss) that have a carbon exchange rate (Cowan & Farquhar, 1977; Buckley *et al.*, 2017b). Assuming a marginal water cost of carbon gain *λ*_w_ [mol H_2_O mol^−1^ CO_2_], the total carbon gain rate per area to maximize is:

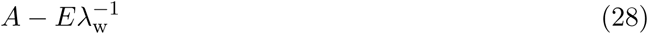

This can be thought of as a profit – carbon gain minus water loss multiplied by a water-to-carbon exchange rate – to be maximized. The optimal solution will be where 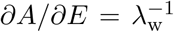. The cost of water increases with the *inverse* of *λ*_w_. For consistency in units, *E* in this equation must be converted from mol to *µ*mol H_2_O m^−2^ s^−1^ during optimization. Traditionally, optimization models find the *g*_sw_ that optimizes carbon gain and water loss, but other traits and other costs can be added for multivariate optimization. Since *SR* also affects carbon gain and water loss, I jointly find the optimum of both stomatal traits, denoted *g*_sw,opt_ and *SR*_opt_. I also included an extrinsic cost of upper stomata (*λ*_*SR*_ [Pa^−1^]) in some models (see below):

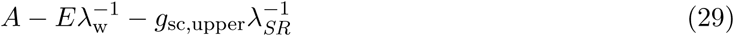

*λ*_*SR*_ must have Pa in the denominator so that 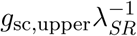 has units *µ*mol CO_2_ m^−2^ s^−1^. The cost of amphistomy is proportional to the *inverse* of *λ*_*SR*_. When 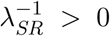, this implies that stomatal conductance through the upper surface incurs some additional cost compared to the same conductance through the lower surface (see Discussion). I refer to 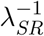 as an ‘extrinsic’ cost of amphistomy because it is an *ad hoc* assumption and not an intrinsic part of the mechanistic model. Since the model does not specify mechanistically how this cost arises, I chose values of 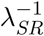 that yielded nontrivial results, but these values are arbitrary and their realism needs to be tested with experiments.

I developed an R package **leafoptimizer** to integrate leaf energy budget models in **tealeaves** and C_3_ photosynthesis models in **photosynthesis** and solve for optimal stomatal traits. **leafoptimizer** takes leaf parameters, environmental parameters, carbon costs, and physical constants as input (Table 1). **leafoptimizer** uses the R package **optimx** (Nash & Varadhan, 2011; Nash, 2014) to numerically solve for the trait optima by iteratively finding 1) the equilibrium *T*_leaf_ then 2) the *E, A*, and net carbon balance (Eq. 29) at that *T*_leaf_ until net carbon balance is maximized. For larger leaves under high light and warm temperatures, *g*_sw,opt_ was often unrealistically high to cool leaves down closer to the optimum for photosynthesis (results not shown). Therefore, I set the maximum *g*_sw,opt_ to 16.43 *µ*mol H_2_O m^−2^ s^−1^ Pa^−1^, equal to *g*_sc_ = 10 *µ*mol CO_2_ m^−2^ s^−1^ Pa^−1^). Following Sharkey *et al.* (2007), I use units for conductance that do not change with with atmospheric pressure because they include Pa in the denominator. Often, conductance is reported in units of mol m^−2^ s^−1^ in the physiological literature. When atmospheric pressure is 100 kPa (which is approximately true near sea level), the nominal conductance in pressure-independent units (*µ*mol m^−2^ s^−1^ Pa^−1^) is 10*×* greater than the value in units of mol m^−2^ s^−1^.

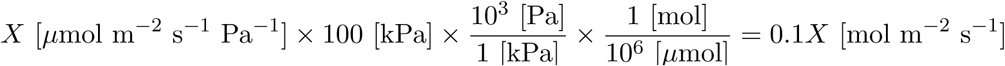

A current version of **leafoptimizer** is available on GitHub (https://github.com/cdmuir/leafoptimizer). The version used for this manuscript (0.0.1) is archived on Zenodo (https://zenodo.org/). I will continue developing the package and depositing revised source code on GitHub between stable release versions. Other scientists can contribute code to improve **leafoptimizer** or modify the source code on their own installations for a more fully customized implementation. A future publication will more fully describe the package and its potential applications. **leafoptimizer** depends on several other R packages: **checkmate** (Lang, 2017), **crayon** (Csárdi, 2017), **dplyr** (Wickham *et al.*, 2018), **glue** (Hester, 2018), **furrr** (Vaughan & Dancho, 2018), **future** (Bengtsson, 2018), **ggplot** (Wickham, 2016), **magrittr** (Bache & Wickham, 2014), **plyr** (Wickham, 2011), **purrr** (Henry & Wickham, 2018a), **rlang** (Henry & Wickham, 2018b), **stringr** (Wickham, 2018), **tibble** (Müller & Wickham, 2019), **tidyr** (Wickham & Henry, 2018), **tidyselect** (Henry & Wickham, 2018c), and **units** (Pebesma *et al.*, 2016).

#### Model 1: no extrinsic cost of amphistomy

Amphistomy increases *E* most at low wind speed and in large leaves (Foster & Smith, 1986, this study), conditions most common in forest understories where amphistomy is rare (Salisbury, 1927; Peat & Fitter, 1994; Muir, 2018). Amphistomy also increases *A* more under high light when CO_2_ limits photosynthesis (Jones, 1985; Mott *et al.*, 1984). Therefore, I hypothesized that the increased cost of *E* and decreased photosynthetic benefit could drive the empirical observation that amphistomy is more common in high light environments (Salisbury, 1927; Mott *et al.*, 1984; Mott & Michaelson, 1991; Peat & Fitter, 1994; Jordan *et al.*, 2014; Bucher *et al.*, 2017; Muir, 2018; Drake *et al.*, 2019). To test whether this hypothesis is plausible, I solved for *g*_sw,opt_ and *SR*_opt_ across a light gradient (PPFD = 100 – 2000) at low (0.2 m s^−1^) and moderate (2 m s^−1^) wind speeds for small (*d* = 0.004 m), medium (*d* = 0.04 m), and large (*d* = 0.4 m) leaves. These values were chosen to ensure that free convection would be important at low wind speeds (see Results). The cost of water was 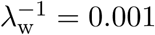 mol CO_2_ mol^−1^ H_2_O. This value is close to that estimated for forbs and grasses under well-watered conditions (0.000981, Manzoni *et al.* (2011)), which is appropriate here because these functional types vary more in stomatal ratio than woody plants (Muir, 2015, 2018) and this study does not evaluate the effects of drought stress, which would increase 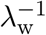. The extrinsic cost of upper stomata was 0. Other model variables and parameters are described in Table S2. Biochemical parameters at 25 ° C for the photosynthesis model roughly match the average and range of values from global plant surveys (Rogers *et al.*, 2017).

#### Model 2: extrinsic cost of amphistomy

A fitness cost of upper stomata would explain the rarity of amphistomy in nature (Metcalfe & Chalk, 1979; Peat & Fitter, 1994; Muir, 2015, 2018; Drake *et al.*, 2019). Model 1 tests whether a cost emerges instrinsically as a result of how stomatal ratio affect *A* and *E*. In this model, I add an exstrinsic cost to upper stomata by varying 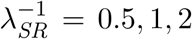 Pa. Higher 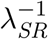 (lower *λ*_*SR*_) corresponds with a higher cost of conductance through upper stomata. Other parameters were the same or similar to Model 1 (Table S3). Because low versus high biochemical parameters *J*_max,25_ and *V*_xmax,25_ had little qualitative effect (see Results), I used a single intermediate value for Models 2 and 3.

#### Model 3: extrinsic cost of amphistomy covaries with light

Covariation between fitness costs and benefits can generate threshold-like clines because there is a very narrow window of environments in which intermediate phenotypes are optimal. I tested this by covarying PPFD and 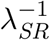, otherwise using the same parameter values as in Model 2 (Table S4). PPFD varied between 73 – 1927. I selected 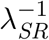 values that weakly, moderately, or strongly covaried with PPFD. 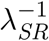 varied the least (0.667 – 1.333) under the weak-covariance scenario and the most (0.002 – 1.998) under the strong-covariance scenario. In all cases, I used bivariate Gaussian covariance stucture, but adjusted the range of 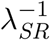, as depicted in Fig. S2.

Source code for these simulations is available on GitHub (https://github.com/cdmuir/stomata-light) and archived on Zenodo (https://zenodo.org/).

## Results

### Amphistomy increases transpiration and CO_2_ assimilation

Output from **tealeaves** and **photosynthesis** packages recapitulate previous work demonstrating that amphistomy increases transpiration (*E*, Fig. 2A) and photosynthetic CO_2_ assimilation (*A*, Fig. 2B). When free convection is important at low wind speed and/or large leaf size, amphistomatous leaves have up to 1.5 times greater *E* than a hypostomatous leaf in the same conditions. The difference in *E* between stomatal ratio phenotypes is less when forced convection prevails at higher wind speeds. Amphistomatous leaves increase photosynthetic rate, all else being equal, by providing an additional parallel pathway for CO_2_ diffusion. Interestingly, leaves with intermediate phenotypes (stomatal ratio [*SR*] = 0.25) increase photosynthetic rate nearly as much as completely amphistomatous leaves (*SR* = 0.5, Fig. 2B).

**Figure 2:**
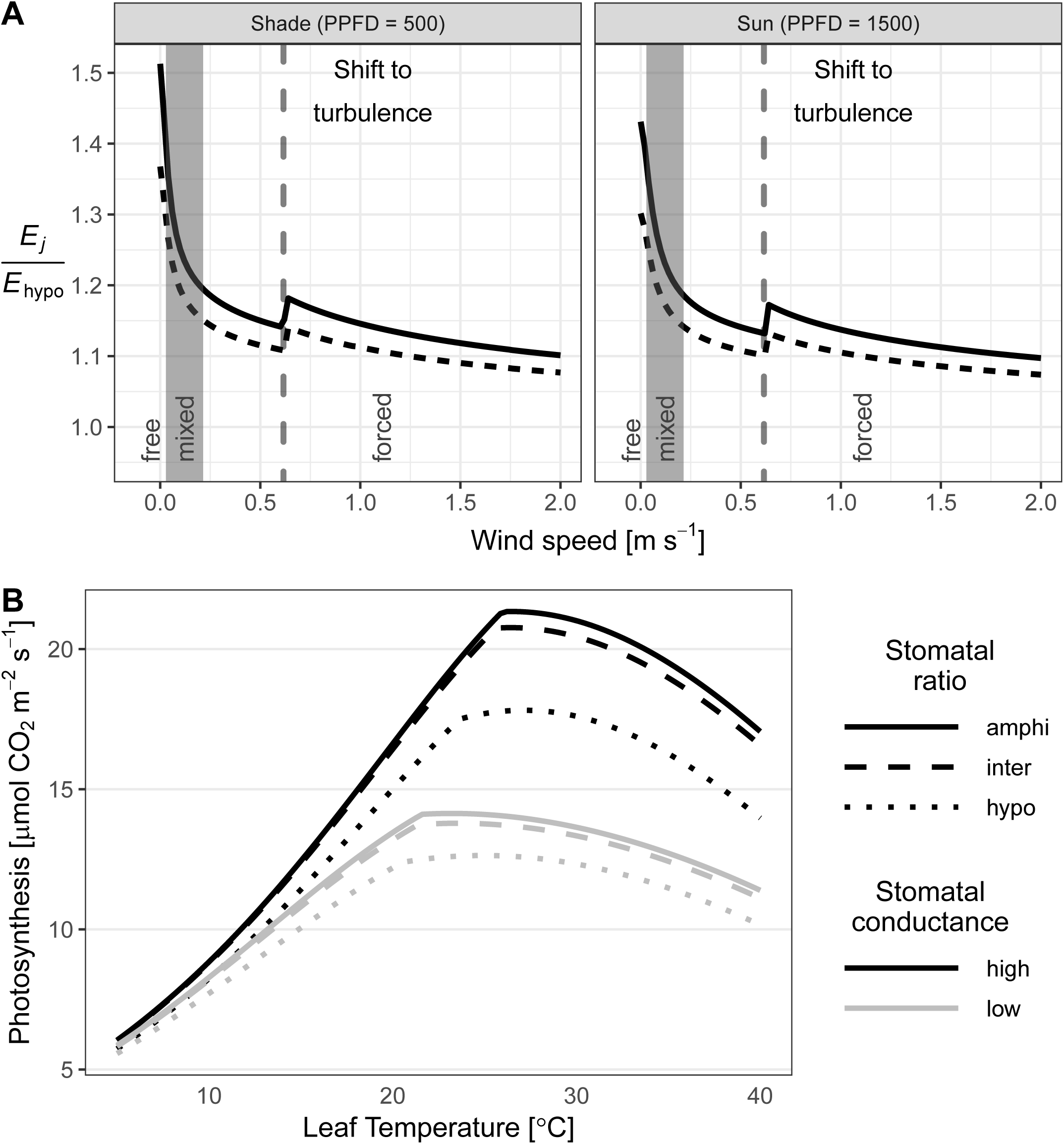
Amphistomy increases transpiration and CO_2_ assimilation. **A**) Output from **tealeaves** shows that amphistomatous (Stomatal Ratio (*SR*) = 0.5, solid black lines) and intermediate (*SR* = 0.25, dashed black lines) leaves transpire more water than hypostomatous leaves (*E*_*j*_*/E*_hypo_ > 1) when stomatal conductance and other leaf/environmental parameters are constant. The effect of *SR* is especially strong at very low wind speeds (*x*-axis) when free convection is significant (low wind speed, *Ar* > 10); it less important for most leaves in which forced convection (high wind speed, *Ar* < 0.1) and turbulent flow (*Re* > 4000, right of dashed line) dominates heat and mass transfer. The effect is similar in both shade (Photosynthetic photon flux density (PPFD) = 500 *µ*mol quanta m^−2^ s^−1^, left facet) and sun (PPFD = 1500 *µ*mol quanta m^−2^ s^−1^, right facet), although total transpiration is greater in the sun (results not shown). **B**). Output from **photosynthesis** shows that amphistomatous leaves (solid lines) increase photosynthetic rate compared to intermediate (dashed lines) and hypostomatous (dotted lines) leaves under the same conditions. The values of *SR* are the same as **A**. Stomatal conductance was set to *g*_sw_ = 1 (low) and 4 (high) *µ*mol H_2_O m^−2^ s^−1^ Pa^−1^. In all conditions, photosynthetic rate peaks at an intermediate temperature. See Materials and Methods for other parameter values.

#### Model 1: Amphistomy is almost always favored when there is no cost of upper stomata

In this model, I used **leafoptimizer** to solve for the *g*_sw,opt_ and *SR*_opt_ that optimally balances *A* and *E* across a range of environmental conditions (Table S2), given a cost of water, but no extrinsic cost of upper stomata.

In almost all areas of parameter space, the additional *A* associated with amphistomy outweighs the increased *E* (Fig. 2). A greater fraction of stomata on the lower surface can be beneficial only when reduced transpiration heats the leaf up closer to the optimum for photosynthesis (*T*_leaf_ ≈ 25°C) given the temperature response parameters assumed in this study [Fig. 2B, Table S1]). This only occurred at suboptimal air temperatures for large leaves in still air at low to moderate irradiance (Fig. 3). Forced convection dominated heat and mass transfer in smaller leaves or leaves in moving air (Figs. 3, S1). Only with the transition to free convection in large leaves and still air does reducing the conductance on the upper surface dramatically decrease transpiration (Fig. 2A). However, this beneficial effect of having lower stomatal conductance on the upper surface goes away under high irradiance because *T*_leaf_ rises toward the optimal temperature for photosynthesis. Hence, amphistomy is always favored at high irradiance when there is no extrinsic cost of upper stomata (Fig. 3). Biochemical parameters had little qualitative effect on the results (Fig. S3).

**Figure 3:**
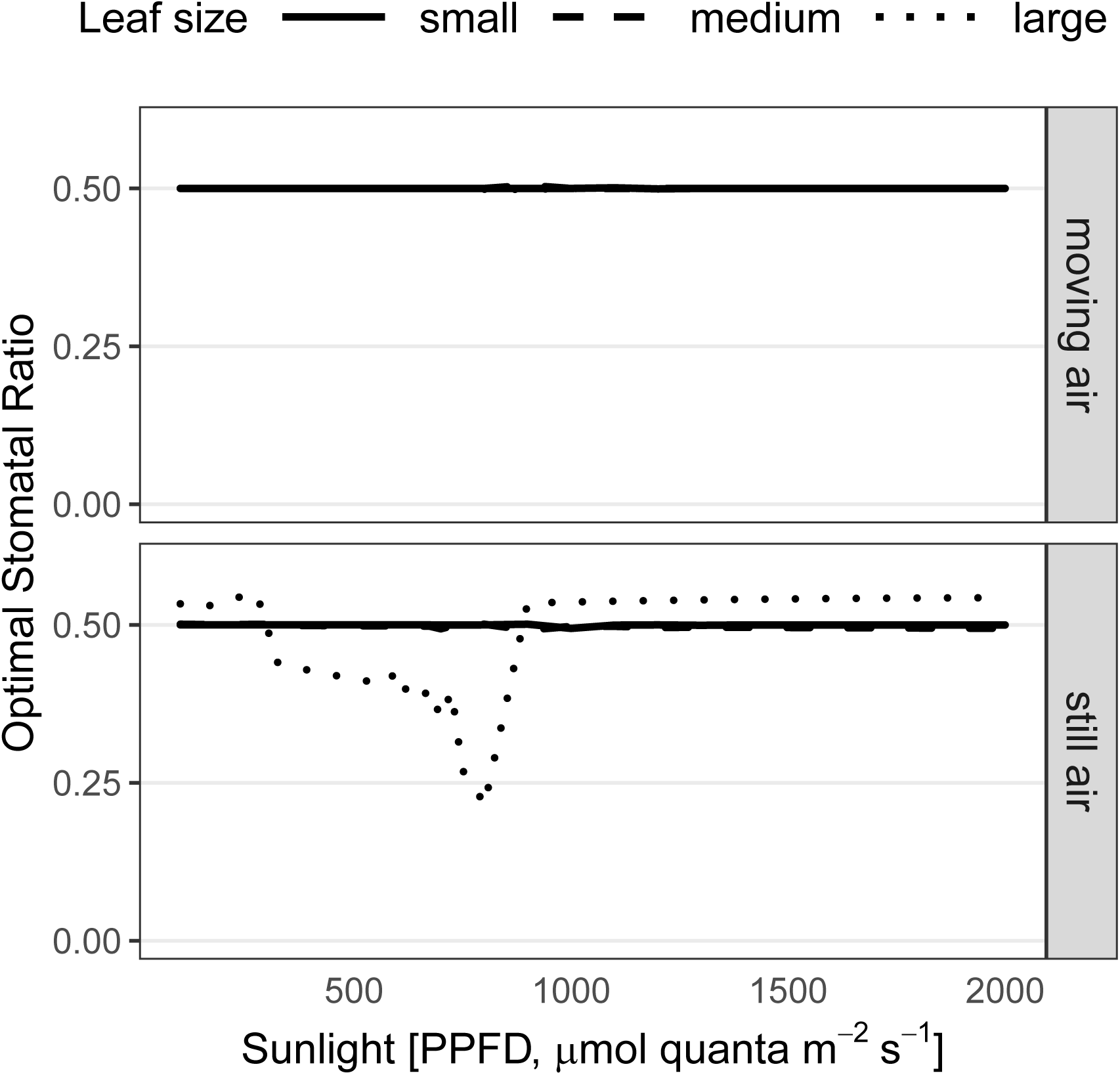
Model 1 shows that amphistomy is almost always optimal when there is no extrinsic cost. The optimal stomatal ratio *SR*_opt_ (*y*-axis) along a PPFD gradient (*x*-axis) for small (*d* = 0.004 m, solid lines), medium (*d* = 0.04 m, dashed lines), and large (*d* = 0.4 m, dotted lines) leaves. In moving air (*u* = 2 m s^−1^, upper facet), amphistomy is always favored; all lines overlap at *SR*_opt_ ≈ 0.5. In still air (*u* = 0.2 m s^−1^, lower facet), *SR*_opt_ < 0.5 only occurs for large leaves in partial shade. Only results for *T*_leaf_ = 25 °C and *J*_max,25_ = 75 shown, but results are qualitatively similar for other variable combinations (Fig. S3). See Table S2 for other parameter values.

#### Model 2: an extrinsic cost of amphistomy produces correlations with light

Model 1 demonstrated that without an extrinsic cost, amphistomy is nearly always optimal. However, under the same leaf and environmental parameters as Model 1, an extrinsic cost leads to many situations in which hypostomy or intermediate *SR* are optimal (Fig. 4A). Under low light, hypostomy is better unless the cost of amphistomy is very low, but under high light, *SR*_opt_ depends strongly on 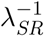. When the cost is low, an intermediate *SR*_opt_ occurs at most light levels; when the cost is high, *SR*_opt_ is always near 0 (hypostomy). This model also predicts some covariation between *SR*_opt_ and *g*_sw,opt_. At low light, both values are predicted to be low; at high light, both values are higher (Fig. 4).

**Figure 4:**
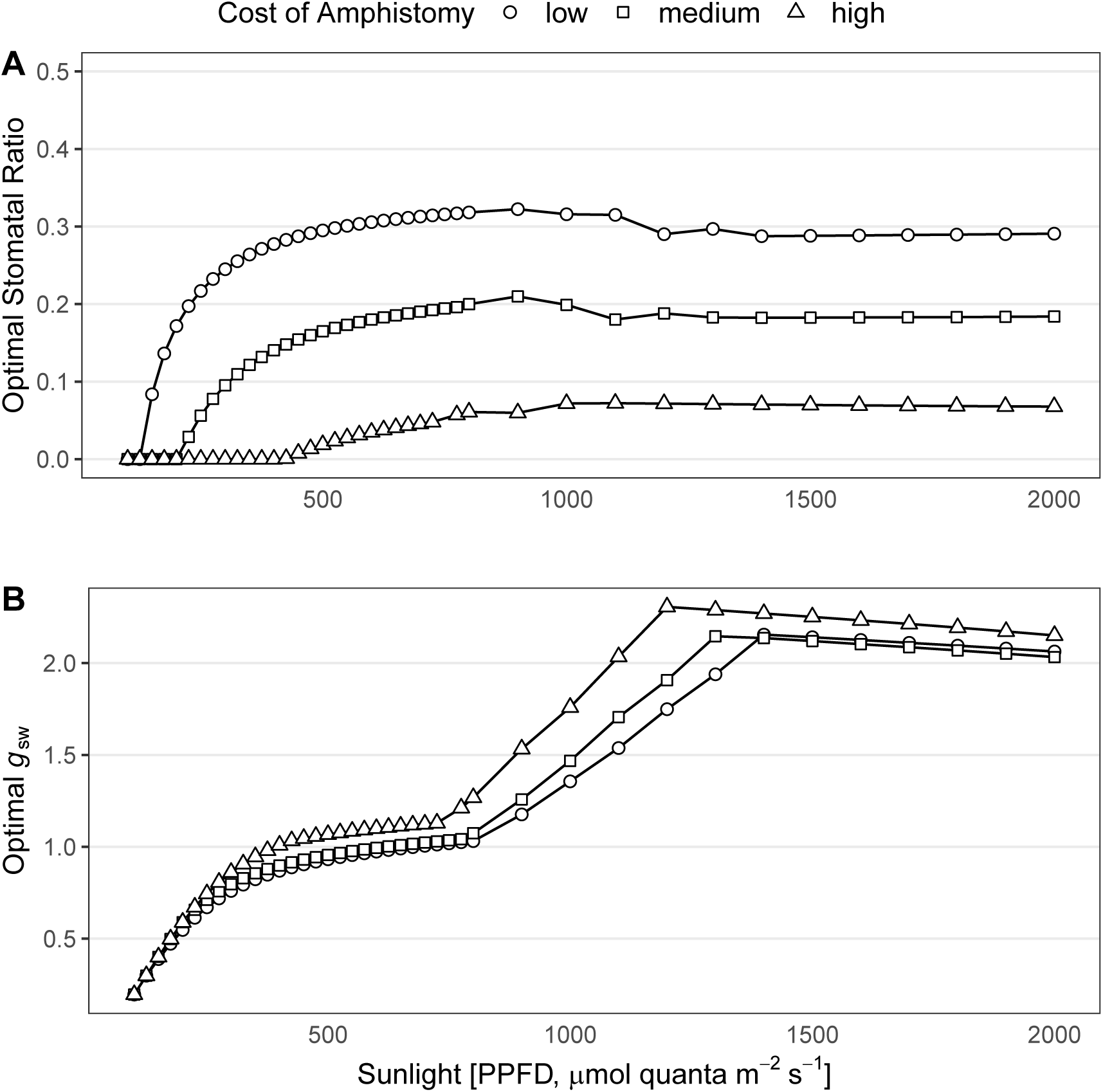
An extrinsic cost of amphistomy generates covariation between sunlight, stomatal ratio, and stomatal conductance. **A**) Model 2 predicts that optimal stomatal ratio (*y*-axis) increases with sunlight (*x*-axis). The optimal value depends on the cost of amphistomy 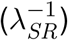: high costs (triangles) favor hypostomy (*SR*_opt_ ≈ 0) over a broad range of light levels; low costs (circles) favor an intermediate value (*SR*_opt_ ≈ 0.3) at most light levels. **B**) Optimal stomatal conductance (*g*_sw_ [*µ*mol H_2_O m^−2^ s^−1^ Pa^−1^], *y*-axis) increases with sunlight, although the pattern is complex. The cost of amphistomy had relatively little effect on the shape of the relationship between sunlight and *g*_sw_, because all three curves follow similar trajectories. See Table S3 for other parameter values.

#### Model 3: low costs of amphistomy at high light can produce threshold-like clines

Compared to Model 2, covariation between costs of amphistomy and light produced stronger threshold-like clines between light and *SR*_opt_ (Fig. 5). With strong covariance, complete hypostomy (*SR*_opt_ = 0) was optimal under low light and high 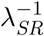; complete amphistomy (*SR*_opt_ = 0.5) was optimal under high light and low 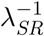. The correlation between *SR*_opt_ and *g*_sw,opt_ was similar to Model 2.

**Figure 5:**
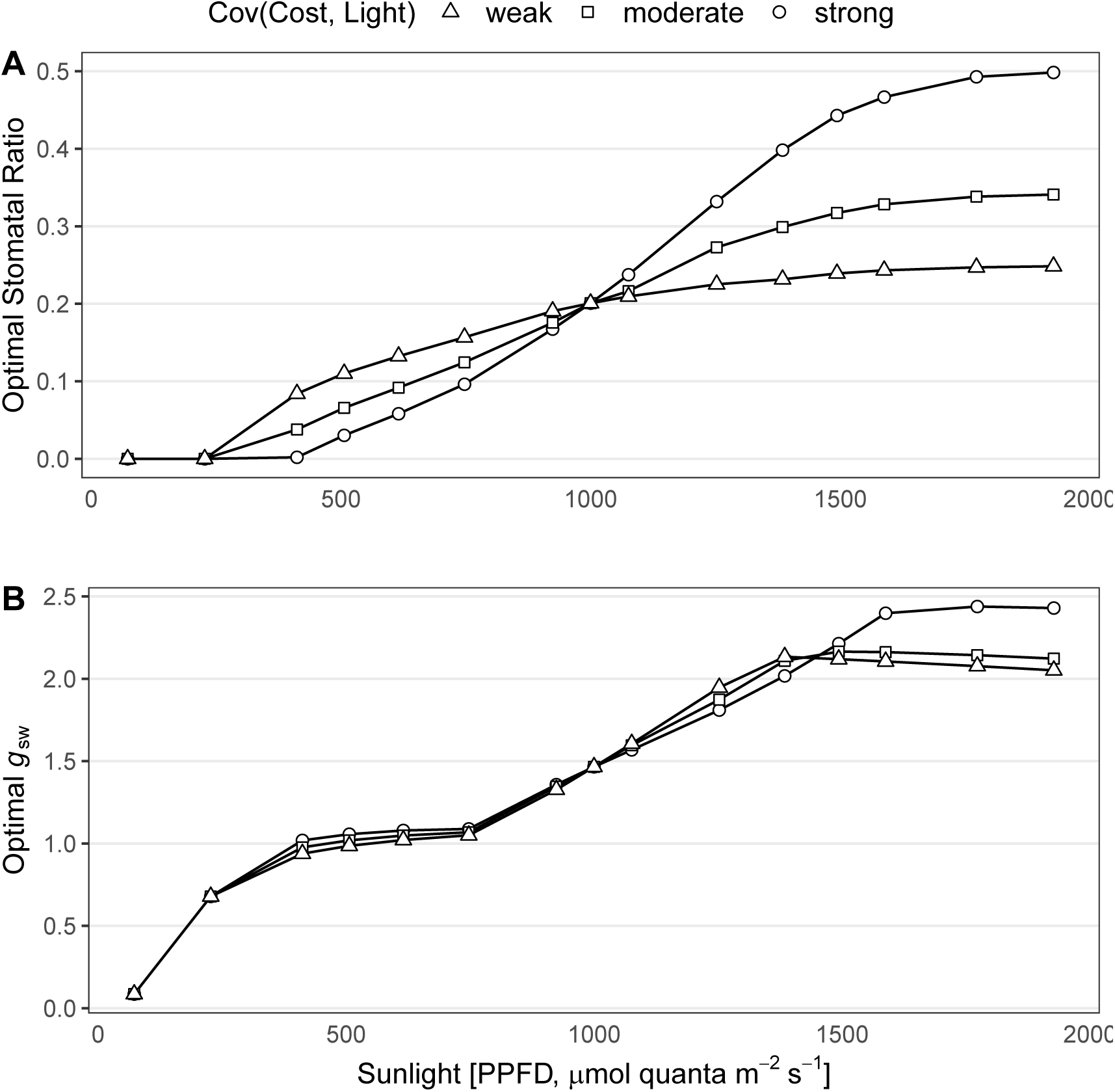
Strong covariance between an extrinsic cost of amphistomy 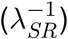 and sunlight [“Cov(Cost, Light)”] generates threshold-like clines between sunlight, stomatal ratio, and stomatal conductance. **A**) Model 3 predicts that optimal stomatal ratio (*y*-axis) increases with sunlight (*x*-axis). When the covariance between costs and light is strong (circles), hypostomy is favored at low light, amphistomy is favored at high light, and there is a nonlinear transition between the two ends. Conversely, when the covariance is low (triangles), intermediate values of *SR*_opt_ are favored at high light, similar to Model 2. **B**) Optimal stomatal conductance (*g*_sw_ [*µ*mol H_2_O m^−2^ s^−1^ Pa^−1^], *y*-axis) increases with sunlight, although the pattern is complex. The covariance between cost and light had relatively little effect on *g*_sw_, because all three curves follow similar trajectories. See Table S4 for other parameter values.

## Discussion

I used three optimality models based on the biophysics and biochemistry of leaf temperature and photosynthesis to predict stomatal ratio (*SR*_opt_) and conductance (*g*_sw,opt_) across light gradients. I draw three substantial conclusions about the evolution of stomatal traits that inform more general questions about phenotypic evolution.

First, a tradeoff between increased photosynthetic CO_2_ assimilation (*A*, 2B) and water loss (*E*, Fig. 2A) does not explain why amphistomy is rare because the benefits almost always outweigh the costs (Model 1, Table 3). Previous modeling and experiments already demonstrated the physiological effects of amphistomy on *A* and *E* (Parkhurst, 1978; Gutschick, 1984; Foster & Smith, 1986; Parkhurst & Mott, 1990; S̆antrůc̆ek *et al.*, 2019), but these insights have not been combined for optimality modeling. Hypostomy is sometimes optimal at low wind speed, low/partial sun, and suboptimal temperatures (Fig. 3, S1) because decreased *E* brings *T*_leaf_ closer to its optimum. However, these restrictive conditions are probably not common in nature; even light wind speeds greater than 1 m s^−1^ would completely eliminate this effect (Fig. 3).

**Table 3:** Which model accodomates empirical observations? The empirical observations are that amphistomy is rare, correlated with light habitat, correlated with stomatal density, and is bimodal. See Introduction for further detail. A ‘✓’ indicates that the model can explain this observation.

| *Amphistomy is:* | Empirical observations |  |  |  |
| --- | --- | --- | --- | --- |
|  | Rare | Cor with light | Cor with stomatal density | Bimodal |
| Model 1: No cost of $SR$ | | | | |
| Model 2: Cost of $SR$ | ✓ | ✓ | ✓ | |
| Model 3: Covarying cost of $SR$ | ✓ | ✓ | ✓ | ✓ |

Second, an extrinsic cost of amphistomy 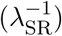 produces a cline between light and *SR*_opt_ (Model 2, Fig. 4). Under the same parameters in Model 1, no such cline is predicted. A previous phenomenological model also suggested that the cost of amphistomy is important (Muir, 2015), but could not distinguish between an “intrinsic” (Model 1) and “extrinsic” (Models 2 and 3) cost. The leaf temperature and photosynthesis models in this study indicate that the tradeoff between *A* and *E* is not the mechanism explaining stomatal ratio, but future mechanistic models of other processes effected by stomatal ratio (e.g. hydraulic conductance outside the xylem (Buckley *et al.*, 2015, 2017a; Drake *et al.*, 2019)) may reveal an ‘intrinsic’ cost. Model 2 also explains why stomatal ratio and conductance positively covary along light gradients (Muir, 2018). Both *SR*_opt_ and *g*_sw,opt_ are beneficial under high light because the marginal benefit of increased CO_2_ supply is greater under high light. I am assuming here that stomatal density is a proxy for operational stomatal conductance (Franks & Beerling, 2009). Generally, stomatal density increases with light up to an intermediate value then decreases slightly (Poorter *et al.*, 2019), consistent with model predictions here (Figs. 4B, 5B). However, in real plants, many other traits change in response to light which are forced to remain constant in the model, so this correspondance between model predictions and experiments may be spurious. Overall, this model indicates that optimizing both density and distribution of stomata on a leaf may help plants fully take advantage of high light and should be considered together in future analyses of light responses.

Third, only when the cost of amphistomy covaries with light does a threshold-like trait-environment relationship emerge (Model 3, Fig. 5). Model 2 explains other empirical observations (Table 3) but fails to explain why intermediate stomatal ratio trait values are rare in nature. Under that model, intermediate values should be common. Only by coupling a benefit of increased *A* under high light with a low cost of amphistomy in the same environment do we predict discrete clusters of hypo- and amphistomatous leaves (i.e. bimodality). Covariation between costs of amphistomy and light may be the only way in this modeling framework to get phenotypic clusters when the underlying environmental gradient is continuous. I used light as an environmental gradient based on *a priori* hypotheses, but covariation between the cost of amphistomy and another environment or trait could produce qualitatively similar results. For example, amphistomy increases *A* more in leaves with high resistance to mesophyll CO_2_ diffusion (Parkhurst, 1978). Covariation between 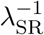 and that trait could also produce a similar effect, but would not necessarily explain why amphistomy is common in high light environments.

The goals of optimality models are to accomodate existing observations and generate new testable predictions. Model 3 accomodates existing observations, but is complex and therefore important to evaluate with future empirical tests of its predictions. In particular, the model implicates the importance of covariation between costs and benefits of amphistomy. Hypostomy is favored in low light with low costs of amphistomy, but high light only favors amphistomy (*SR*_opt_ = 0.5) when costs are also low. This is important because some proposed costs probably do not covary with light gradients this way, while others likely do. For example, amphistomy can dehydrate the palisade mesophyll when there is strong evaporative demand (Buckley *et al.*, 2017a; Drake *et al.*, 2019), but this cost should be stronger, not weaker, under high light. Furthermore, when leaves can optimize both *SR* and *g*_sw_ simultaneously, amphistomatous leaves have lower *g*_sw,opt_ and hence lower evaporative demand than hypostomatous plants holding all else constant (Fig. 4). Amphistomy may also be costly if it increases susceptibility to foliar pathogens that are more likely to land on the upper surface of a horizontally oriented leaf (Gutschick, 1984; McKown *et al.*, 2014). Because many pathogens need a wet leaf microclimate to germinate and grow, a leaf in high light that dries faster is less likely to experience this cost than one in the shade. Hence, if pathogens are the primary cost of amphistomy, then this cost should be higher in shady habitats and lower in sunny habitats, consistent with the assumptions of Model 3. Future work should focus on identifying the abiotic and biotic cost(s) of upper stomata at different light levels under natural conditions. We also need to evaluate how often the distribution of light values is unimodal in nature (hypothesis 2) and the role of developmental constraints on stomatal evolution (hypothesis 1).

There are several important limitations of this study that will need to be addressed in future work. Currently **leafoptimizer** only optimizes stomatal traits while other traits are held constant. But traits such as leaf size, mesophyll conductance, *J*_max_/*V*_cmax_ acclimate and evolve too. If all these traits could vary together in the model, different patterns might emerge. For example, high light favors thick leaves to capture more photons and greater investment in photosynthetic biochemistry, traits that make increased CO_2_ supply more advantageous. In this case, a greater benefit rather than increased cost might explain why amphistomy is common at high light. Furthermore, this study did not exhaustively explore relevant parameter space. It is possible that further exploration may reveal patterns not identified here. For example, I only used a single set of temperature response functions, even those these vary within and between species (Medlyn *et al.*, 2002; Slot & Winter, 2017). However, this limitation does not qualitatively change the result that amphistomy only significantly affects evaporation, and hence leaf temperature, when leaf size is large, wind speed is almost zero, and there is relatively high sunlight. These conditions are not common in nature. Different temperature response parameters that change optimum leaf temperature would alter the range of air temperatures in which hypostomy would helps keep leaf temperature closer its optimum under restrictive parameter spece. The model also uses bulk leaf properties of temperature and photosynthesis at one time point, ignoring spatial variation within the leaf and temporal variation in the environment, which might yield different predictions (Buckley *et al.*, 2017a; Earles *et al.*, 2019). Finally, carbon gain and water loss are not fitness, which is what natural selection cares about. Future theoretical and empirical studies should integrate plant survivorship and reproduction with stomatal function.

Amphistomy is rare despite the fact that it increases photosynthetic rate. Why? Optimality models show this is not because the increased carbon gain is offset by additional water loss. Instead, an additional cost of amphistomy, yet to be identified, must explain why it is rare. Optimality models also predict that amphistomy is common in high light habitats not just because it increases carbon gain but also because the costs of amphistomy are lower. Covariation between costs and benefits may also explain why stomatal ratio forms discrete phenotypic clusters.

## Acknowledgements

I would like to thank Joseph Stinziano and an anonymous reviewer for valuable feedback on this work.

## Supporting Information

### Photosynthetic temperature responses

I calcualted *g*_mc_, Γ^*^, *J*_max_, *K*_C_, *K*_O_, *R*_d_, *V*_cmax_, and *V*_tpu_ at *T*_leaf_ (Table 2) based on an assumed value at 25 °C (Table 1) and temperature response paramters from (Bernacchi *et al.*, 2002, Table S1). Parameters with an exponentially increasing response to temperature were modeled as:

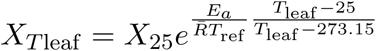

and those with a humped-shaped response were modeled as:

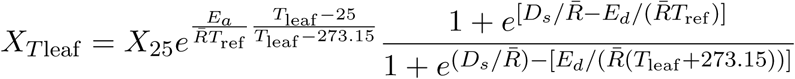

*E*_*a*_ and *E*_*d*_ are the enthalpies of activation and deactivation, respectively, and *D*_*s*_ is the entropy. *T*_ref_ is a reference temperature (25 °C) in K; *T*_leaf_ is a reference temperature in °C.

**Table S1:**
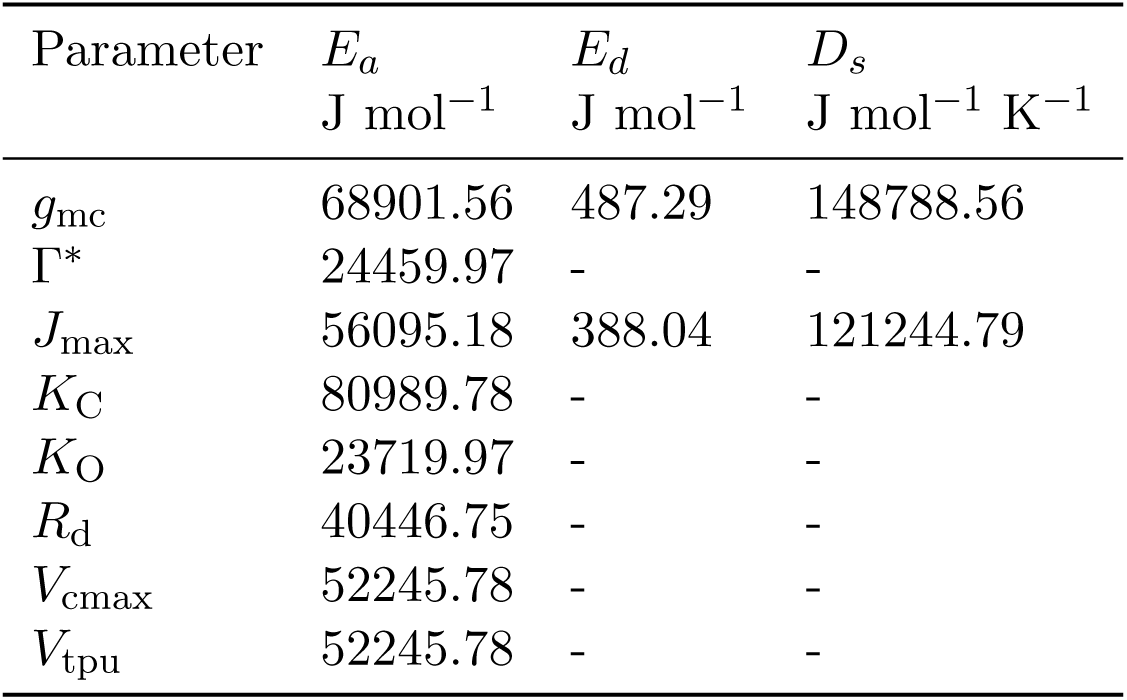
Temperature response parameters

### Parameter conversions in leafoptimizer

Because of their differing origins and uses, leaf energy budget and photosynthesis models sometimes employ different units for the same parameter. As standalone packages, **tealeaves** and **photosynthesis** honor these conventions, but **leafoptimizer** must convert between them. Here I document these conversions.

As noted in the Materials and Methods section, conductance values are converted from m s^−1^ (**tealeaves**) to *µ*mol m^−2^ s^−1^ Pa^−1^ (**photosynthesis**) using the ideal gas law:

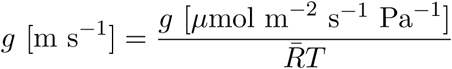

Conductance to water vapor and CO_2_ are interconverted using the the `gc2gw()` and `gw2qc()` functions:

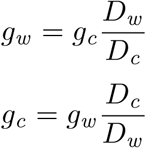

Incident shortwave radiation (*S*_sw_ [W m^−2^], **tealeaves**) is interconverted with PPFD [*µ*mol quanta m^−2^ s^−1^] (**photosynthesis**) following Gutschick (2016) using the functions `sun2ppfd()` and `ppfd2sun()`. Shortwave radiation is (at first approximation) the sum of photosynthetically active radiation (PAR) and near-infrared radiation (NIR):

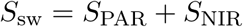

Most sources (e.g. Jones, 2014) assume that *S*_PAR_ = *S*_NIR_ for sunlight, so *f*_PAR_ = 0.5. To convert PAR to PPFD, divide by the energy per mol quanta. assuming *E*_*q*_ = 220 kJ mol^−1^ quanta for PAR:

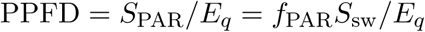

**tealeaves** uses stomatal ratio (*SR*), while **photosynthesis** uses a partitioning factor *k*_sc_. These are automatically interconverted as:

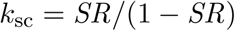

---

**Figure S1:**
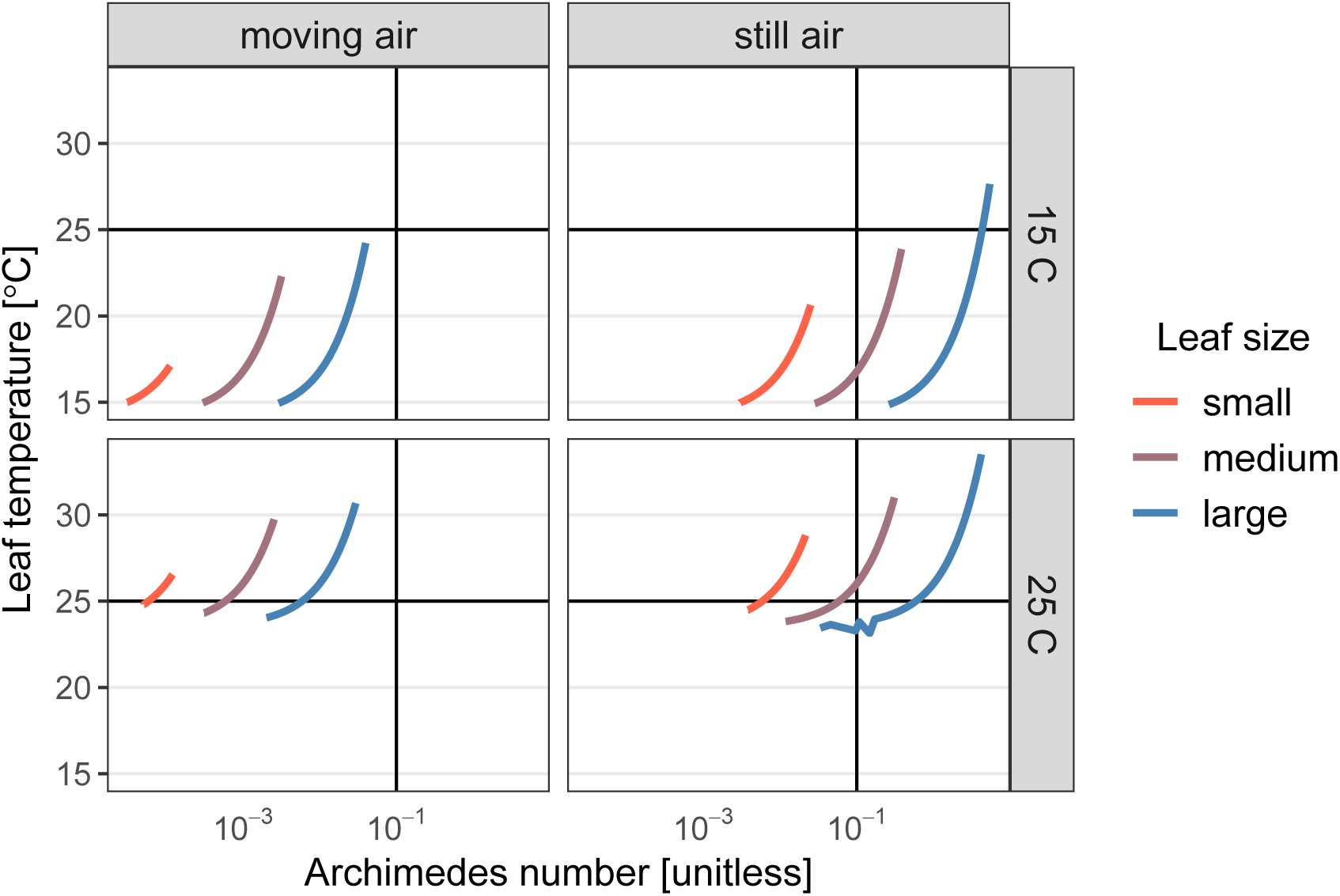
Environmental conditions that favor hypostomy are rare when there is no extrinsic cost to amphistomy. Each facet plots the Archimedes number (*x*-axis) against leaf temperature (*y*-axis) in moving air (*u* = 2 m s^−1^, left column) and still air (*u* = 2 m s^−1^, right column) at two air temperatures: *T*_air_ = 15 °C (top row) and *T*_air_ = 25 °C, (bottom row). Results from a third temperature (*T*_air_ = 35 °C) indicated bistability at intermediate Archimedes numbers and are not shown here, but results are archived online (see Materials and Methods). Hypostomy reduces transpiration, increasing leaf temperature, which can be beneficial when leaf temperatures are suboptimal for photosynthesis (approximately 25 °C, horizontal line in all facets). This only occurs at low air temperatures (top row). Furthermore, free convection must be significant (Archimedes number > 0.1, vertical line in all facets). This only occurs for medium and large leaves under high light, which generates a larger leaf-to-air temperature differential. See Table S2 for other parameter values.

**Figure S2:**
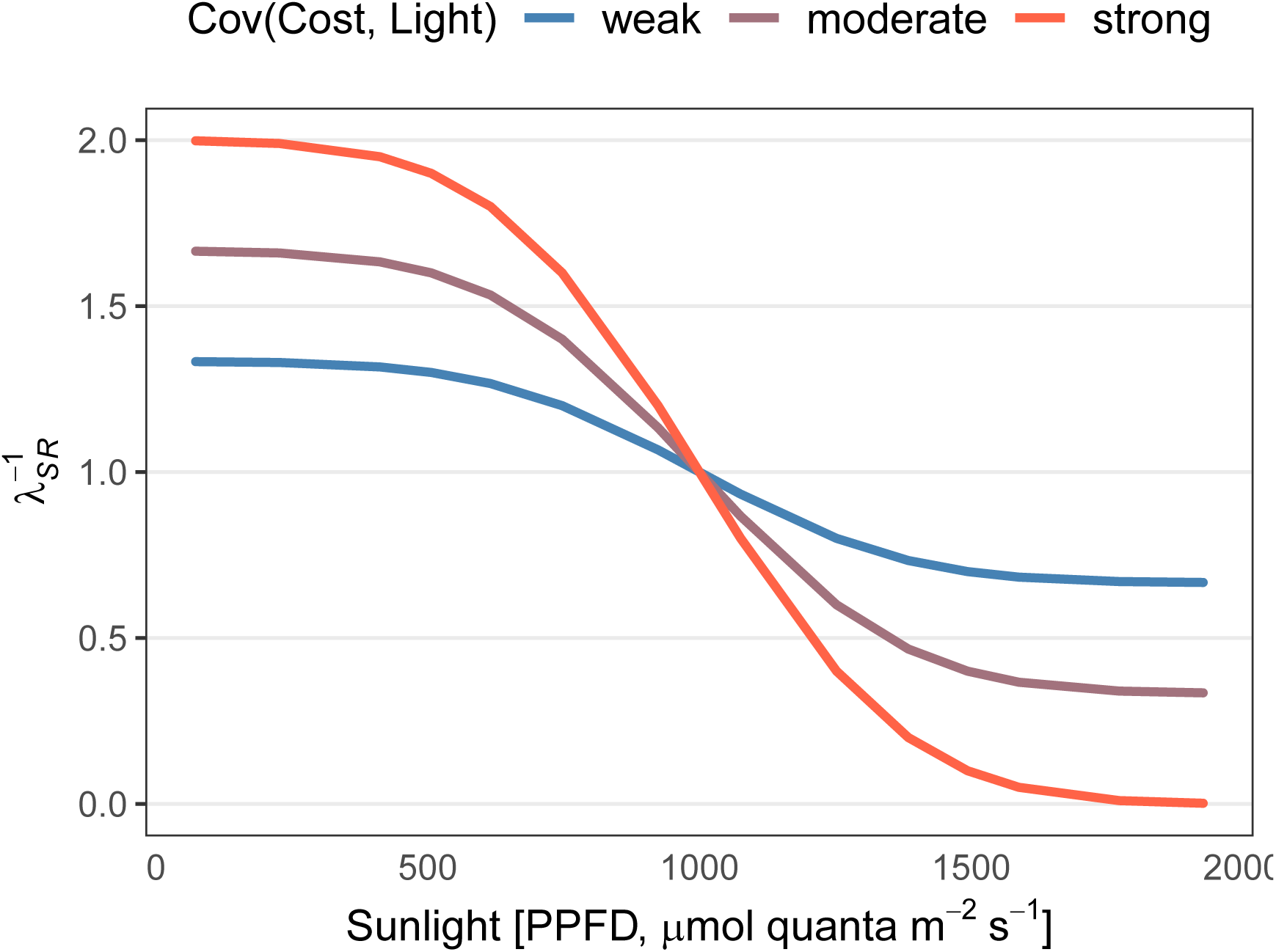
Weak, moderate, and strong examples of covariation [“Cov(Cost, Light)”] between light (PPFD, *x*-axis) and the cost of amphistomy (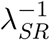 Pa, *y*-axis) used in Model 3. The inverse of *λ*_*SR*_ is plotted because as this value increases, the costs of amphistomy are greater. When the covariance is strong, 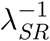 has greater range over the same PPFD values compared to when the covariance is weak.

**Figure S3:**
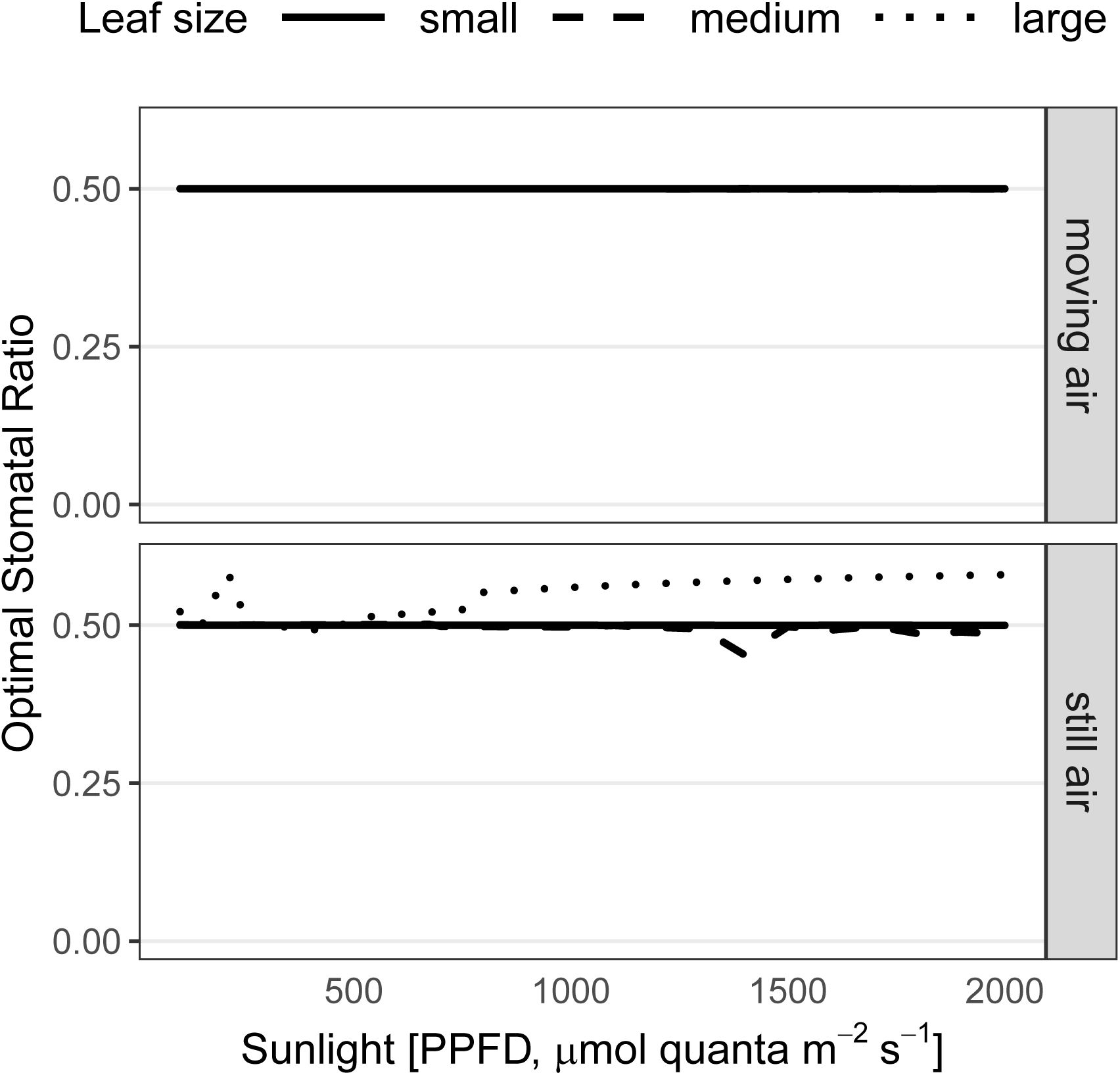
Model 1 shows that amphistomy is almost always optimal when there is no extrinsic cost, regardless of biochemical parameters. This figure is same as Fig. 3, except the biochemical parameters. The optimal stomatal ratio *SR*_opt_ (*y*-axis) along a PPFD gradient (*x*-axis) for small (*d* = 0.004 m, solid lines), medium (*d* = 0.04 m, dashed lines), and large (*d* = 0.4 m, dotted lines) leaves. In moving air (*u* = 2 m s^−1^, upper facet), amphistomy is always favored; all lines overlap at *SR*_opt_ ≈ 0.5. Only results for *T*_leaf_ = 25 °C and *J*_max,25_ = 150 shown. See Table S2 for other parameter values.

**Table S2:** Model 1 variable and parameter values. See Table 1 for symbol definitions and values of physical constants.

| Symbol | Value(s) | Units |
| --- | --- | --- |
| **Variables:** |  |  |
| $d$ | 0.004, 0.04, 0.4 | m |
| $J_{\max,25}$ | 75, 150 | $\mu\text{mol CO}_2 \text{ m}^{-2} \text{ s}^{-1}$ |
| PPFD | 100 – 2000 | $\mu\text{mol quanta m}^{-2} \text{ s}^{-1}$ |
| $u$ | 0.2, 2 | $\text{m s}^{-1}$ |
| $V_{\text{cmax},25}$ | $2/3 J_{\max,25}$ | $\mu\text{mol CO}_2 \text{ m}^{-2} \text{ s}^{-1}$ |
| **Carbon costs:** |  |  |
| $\lambda_{\text{w}}^{-1}$ | 0.001 | $\text{mol CO}_2 \text{ mol}^{-1} \text{ H}_2\text{O}$ |
| $\lambda_{\text{SR}}^{-1}$ | 0 | Pa |
| **Fixed leaf parameters:** |  |  |
| $\alpha_{\text{s}}$ | 0.5 | none |
| $\alpha_{\text{l}}$ | 0.97 | none |
| $g_{\text{mc},25}$ | 3 | $\mu\text{mol CO}_2 \text{ m}^{-2} \text{ s}^{-1} \text{ Pa}^{-1}$ |
| $g_{\text{uc}}$ | 0.1 | $\mu\text{mol CO}_2 \text{ m}^{-2} \text{ s}^{-1} \text{ Pa}^{-1}$ |
| $\Gamma_{25}^*$ | 3.743 | Pa |
| $K_{\text{C},25}$ | 27.238 | Pa |
| $K_{\text{O},25}$ | 16.582 | kPa |
| $k_{\text{mc}}$ | 1 | none |
| $k_{\text{uc}}$ | 1 | none |
| $\phi_J$ | 0.331 | none |
| $R_{\text{d},25}$ | 2 | $\mu\text{mol CO}_2 \text{ m}^{-2} \text{ s}^{-1}$ |
| $\theta_J$ | 0.825 | none |
| $V_{\text{tpu},25}$ | 200 | $\mu\text{mol CO}_2 \text{ m}^{-2} \text{ s}^{-1}$ |
| **Fixed environmental parameters:** |  |  |
| $C_{\text{air}}$ | 41 | Pa |
| $E_{\text{q}}$ | 220 | $\text{kJ mol}^{-1}$ |
| $f_{\text{PAR}}$ | 0.5 | none |
| $O$ | 21.27565 | kPa |
| $P$ | 101.3246 | kPa |
| $r$ | 0.2 | none |
| $RH$ | 0.5 | none |

**Table S3:** Model 2 variable and parameter values. See Table 1 for symbol definitions and values of physical constants.

| Symbol | Value(s) | Units |
| --- | --- | --- |
| **Variables:** |  |  |
| PPFD | 100 – 2000 | $\mu\text{mol quanta m}^{-2} \text{ s}^{-1}$ |
| **Carbon costs:** |  |  |
| $\lambda_w^{-1}$ | 0.001 | $\text{mol CO}_2 \text{ mol}^{-1} \text{ H}_2\text{O}$ |
| $\lambda_{SR}^{-1}$ | 0.5, 1, 2 | Pa |
| **Fixed leaf parameters:** |  |  |
| $\alpha_s$ | 0.5 | none |
| $\alpha_l$ | 0.97 | none |
| $d$ | 0.1 | m |
| $J_{\text{max},25}$ | 112.5 | $\mu\text{mol CO}_2 \text{ m}^{-2} \text{ s}^{-1}$ |
| $g_{\text{mc},25}$ | 3 | $\mu\text{mol CO}_2 \text{ m}^{-2} \text{ s}^{-1} \text{ Pa}^{-1}$ |
| $g_{\text{uc}}$ | 0.1 | $\mu\text{mol CO}_2 \text{ m}^{-2} \text{ s}^{-1} \text{ Pa}^{-1}$ |
| $\Gamma_{25}^*$ | 3.743 | Pa |
| $K_{\text{C},25}$ | 27.238 | Pa |
| $K_{\text{O},25}$ | 16.582 | kPa |
| $k_{\text{mc}}$ | 1 | none |
| $k_{\text{uc}}$ | 1 | none |
| $\phi_J$ | 0.331 | none |
| $R_{\text{d},25}$ | 2 | $\mu\text{mol CO}_2 \text{ m}^{-2} \text{ s}^{-1}$ |
| $\theta_J$ | 0.825 | none |
| $u$ | 2 | $\text{m s}^{-1}$ |
| $V_{\text{cmax},25}$ | $2/3 J_{\text{max},25}$ | $\mu\text{mol CO}_2 \text{ m}^{-2} \text{ s}^{-1}$ |
| $V_{\text{tpu},25}$ | 200 | $\mu\text{mol CO}_2 \text{ m}^{-2} \text{ s}^{-1}$ |
| **Fixed environmental parameters:** |  |  |
| $C_{\text{air}}$ | 41 | Pa |
| $E_q$ | 220 | $\text{kJ mol}^{-1}$ |
| $f_{\text{PAR}}$ | 0.5 | none |
| $O$ | 21.27565 | kPa |
| $P$ | 101.3246 | kPa |
| $r$ | 0.2 | none |
| $RH$ | 0.5 | none |

**Table S4:**
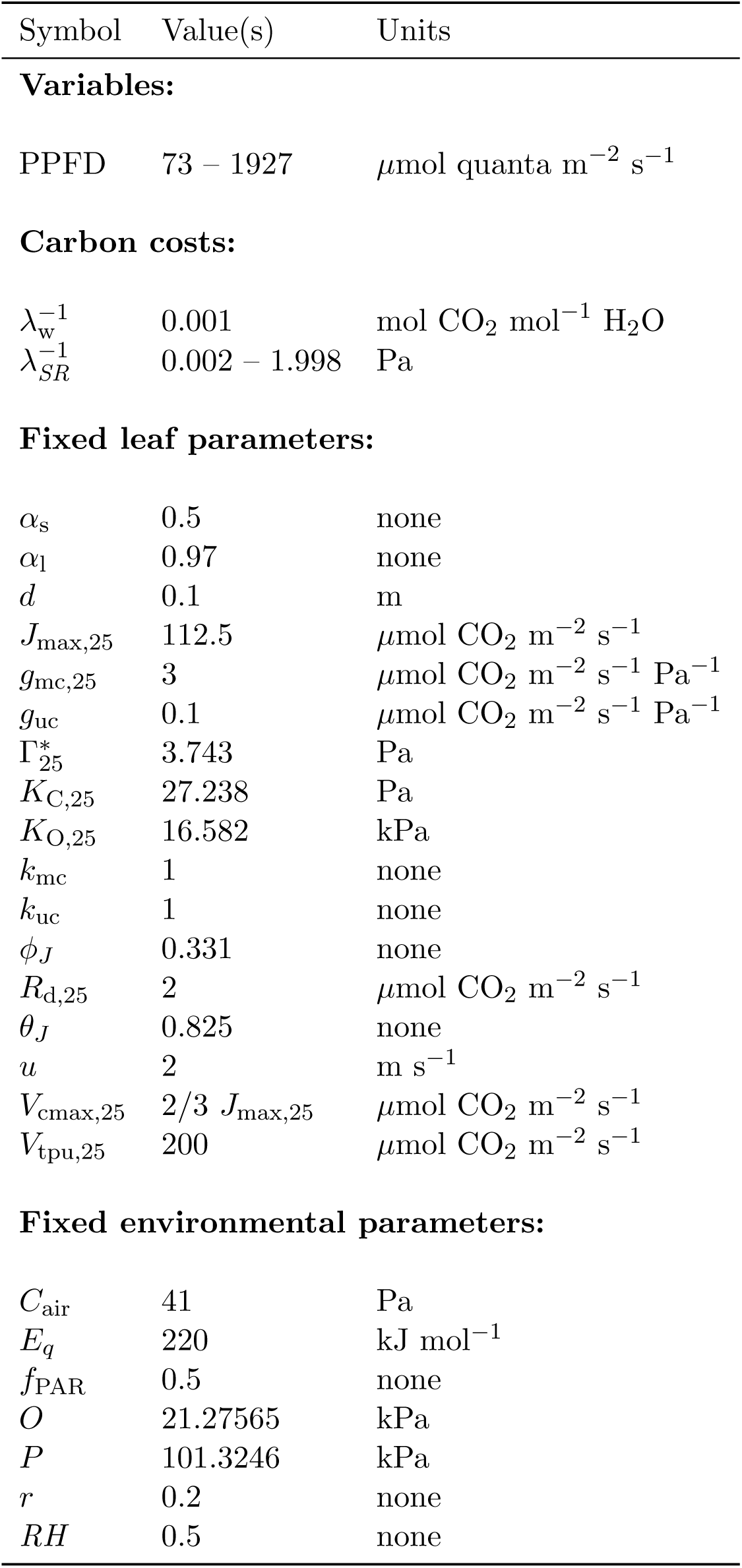
Model 3 variable and parameter values. See Table 1 for symbol definitions and values of physical constants.

| Symbol | Value(s) | Units |
| --- | --- | --- |
| **Variables:** |  |  |
| PPFD | 73 – 1927 | $\mu\text{mol quanta m}^{-2} \text{ s}^{-1}$ |
| **Carbon costs:** |  |  |
| $\lambda_w^{-1}$ | 0.001 | $\text{mol CO}_2 \text{ mol}^{-1} \text{ H}_2\text{O}$ |
| $\lambda_{SR}^{-1}$ | 0.002 – 1.998 | Pa |
| **Fixed leaf parameters:** |  |  |
| $\alpha_s$ | 0.5 | none |
| $\alpha_l$ | 0.97 | none |
| $d$ | 0.1 | m |
| $J_{\text{max},25}$ | 112.5 | $\mu\text{mol CO}_2 \text{ m}^{-2} \text{ s}^{-1}$ |
| $g_{\text{mc},25}$ | 3 | $\mu\text{mol CO}_2 \text{ m}^{-2} \text{ s}^{-1} \text{ Pa}^{-1}$ |
| $g_{\text{uc}}$ | 0.1 | $\mu\text{mol CO}_2 \text{ m}^{-2} \text{ s}^{-1} \text{ Pa}^{-1}$ |
| $\Gamma_{25}^*$ | 3.743 | Pa |
| $K_{\text{C},25}$ | 27.238 | Pa |
| $K_{\text{O},25}$ | 16.582 | kPa |
| $k_{\text{mc}}$ | 1 | none |
| $k_{\text{uc}}$ | 1 | none |
| $\phi_J$ | 0.331 | none |
| $R_{\text{d},25}$ | 2 | $\mu\text{mol CO}_2 \text{ m}^{-2} \text{ s}^{-1}$ |
| $\theta_J$ | 0.825 | none |
| $u$ | 2 | $\text{m s}^{-1}$ |
| $V_{\text{cmax},25}$ | $2/3 J_{\text{max},25}$ | $\mu\text{mol CO}_2 \text{ m}^{-2} \text{ s}^{-1}$ |
| $V_{\text{tpu},25}$ | 200 | $\mu\text{mol CO}_2 \text{ m}^{-2} \text{ s}^{-1}$ |
| **Fixed environmental parameters:** |  |  |
| $C_{\text{air}}$ | 41 | Pa |
| $E_q$ | 220 | $\text{kJ mol}^{-1}$ |
| $f_{\text{PAR}}$ | 0.5 | none |
| $O$ | 21.27565 | kPa |
| $P$ | 101.3246 | kPa |
| $r$ | 0.2 | none |
| $RH$ | 0.5 | none |

